# Thalamo-insular pathway regulates tic generation via motor-limbic crosstalk

**DOI:** 10.1101/2025.08.24.672034

**Authors:** Hiroto Kuno, Natsumi Tsuji, Kenta Kobayashi, Toru Takumi, Yoshihisa Tachibana

**Author notes:** **Corresponding author:** Dr. Yoshihisa Tachibana Department of Physiology and Cell Biology, Kobe University Graduate School of Medicine, Kobe, Japan. 7-5-1 Kusunoki-cho, Chuo-ku, Kobe, Hyogo 650-0017, Japan.

## Abstract

Tic disorders accompanied by premonitory urges are hallmark symptoms of Tourette syndrome (TS), yet the underlying neuronal mechanisms remain elusive. Here, we establish a mouse model of motor tics by unilateral striatal injection of a GABA_A_ receptor antagonist. This model induces c-Fos activation in both motor and limbic structures, including the insular cortex (IC). Fiber photometry reveals tic-associated activity in IC as well as the primary motor cortex (M1). Viral tracing demonstrates that basal ganglia outputs from the substantia nigra pars reticulata are transmitted to IC via the intralaminar thalamic nuclei (ITN). Chemogenetic inhibition of IC or the thalamo-insular pathway suppresses tic-related cortical activity and alleviates tic-like behaviors. These findings identify IC as a key node in tic generation and highlight ITN as critical relay stations linking motor and limbic circuits. Aberrant thalamo-insular signaling thus contributes to the pathophysiology of tic disorders and represents a potential therapeutic target in TS.

**Highlights:**

1. c-Fos activation occurs in both motor and limbic structures in a mouse tic model
2. Basal ganglia outputs are transmitted to insula via intralaminar thalamic nuclei
3. Chemogenetic inhibition of thalamo-insular pathway suppresses motor tics
4. Thalamo-insular inhibition diminishes tic-associated activity in primary motor cortex

**In brief:** Abnormal basal ganglia activity originating from the striatum drives dysfunction of the insular cortex via the intralaminar thalamic nuclei, leading to motor tics. Chemogenetic inhibition of this pathway suppresses tic-like behaviors, highlighting motor-limbic circuit pathology in tic disorders.

## INTRODUCTION

Motor and vocal tics are characterized by sudden, rapid, repetitive movements or involuntary vocalizations.^1^ These symptoms are often preceded by an uncomfortable sensation, known as a premonitory urge, which can be transiently relieved by tic expression.^2–5^ Gilles de la Tourette syndrome (TS) is diagnosed by the presence of multiple motor tics and at least one vocal tic persisting for over a year,^1^ and is frequently accompanied by comorbidities such as obsessive-compulsive disorder (OCD) and attention-deficit/hyperactivity disorder (ADHD).^6–8^ TS typically begins in childhood, affecting approximately 0.3–1.0% of school-aged children.^1,9–12^ While symptoms often improve during adolescence, they can persist or even worsen into adulthood.^1,6^

Current treatment strategies for TS patients include behavioral therapy, pharmacological agents, and, in severe cases, surgical interventions.^13^ Despite extensive clinical characterization, however, the neuronal mechanisms driving tic generation remain incompletely understood. Accumulating evidence implicates dysfunction of the cortico-basal ganglia (BG)-thalamo-cortical (CBGTC) circuits in the pathophysiology of TS.^14^ Microinjections of GABA_A_ receptor antagonists into the sensorimotor striatum reliably induce motor tics in rats and monkeys.^15–19^ Notably, similar manipulations in the associative or limbic subdivisions of the monkey striatum produce distinct behavioral phenotypes: hyperactive and impulsive behaviors resembling ADHD symptoms, or stereotyped behaviors reminiscent of OCD, respectively.^19^ These findings support the “striatal disinhibition hypothesis”, which posits that local GABAergic deficits underlie the pathophysiology of TS symptoms. The striatum contains diverse subtypes of GABAergic interneurons: parvalbumin (PV)-, calretinin-, and neuropeptide Y-positive neurons.^20^ Postmortem studies have revealed a selective reduction of striatal GABAergic interneurons in TS patients.^21,22^ In parallel, the “hyperactive dopamine hypothesis” also provides a compelling framework since pharmacological interventions with dopamine receptor antagonists (i.e., antipsychotics) are clinically effective in suppressing tic symptoms in TS patients.^6,13,23^ However, intrastriatal administration of dopamine agonists in rodents induces stereotyped behaviors, rather than motor tics.^24^ Taken together, these findings highlight that striatal disinhibition-based animal models provide a mechanistically relevant approach for investigating the neuronal circuits underlying tic generation.

Here, we report that striatal injection of bicuculline, a GABA_A_ receptor antagonist, induces motor tics in mice. This model induces activation in both motor and limbic structures, including the insular cortex (IC). We identify an output pathway from BG to IC via the intralaminar thalamic nuclei (ITN), and demonstrate that chemogenetic inhibition of this pathway alleviates tic-like behaviors. These results reveal a motor-limbic circuit that contributes to tic generation.

## RESULTS

### Unilateral striatal bicuculline injection induces tic-like movements

We established a mouse model of motor tics by unilateral injection of bicuculline, a GABA_A_ receptor antagonist, at a low dose (0.2 µg/µl; 0.20 µl per injection) into the motor domain of the dorsal striatum (dSTR) in head-fixed mice (Figure 1). This paradigm recapitulates striatal disinhibition protocols (Figure S1A) previously shown to induce motor tics in rats and monkeys.^15–19^ To monitor the tic-associated cortical activity, we simultaneously recorded electroencephalogram (EEG) signals from the primary motor cortex (M1) ipsilateral to the injection site, together with electromyogram (EMG) activity from the contralateral triceps muscle (Figure 1A). Bicuculline injection into the dSTR produced somatotopically organized tic-like movements in contralateral body parts (Figure S1B and Video S1), depending on the dorsoventral striatal level, as previously reported.^15,16^ In this study, we targeted the striatal injection site (AP: 0.0 mm, ML: 2.7 mm, DV: 3.5 mm from the bregma; Figure 1B), which reliably induced tic-like movements mainly in the contralateral forelimb and orofacial regions (Figure S1B and Video S1). EEG spikes in M1 preceded EMG bursts and kinematic acceleration of the corresponding body parts (Figures 1C-1E for the wrist; S1C-S1E for the mouth). On average, EEG spikes preceded EMG bursts by 17.0 ± 3.7 ms and movement acceleration by 20.7 ± 2.5 ms (mean ± SD).

**Figure 1.**
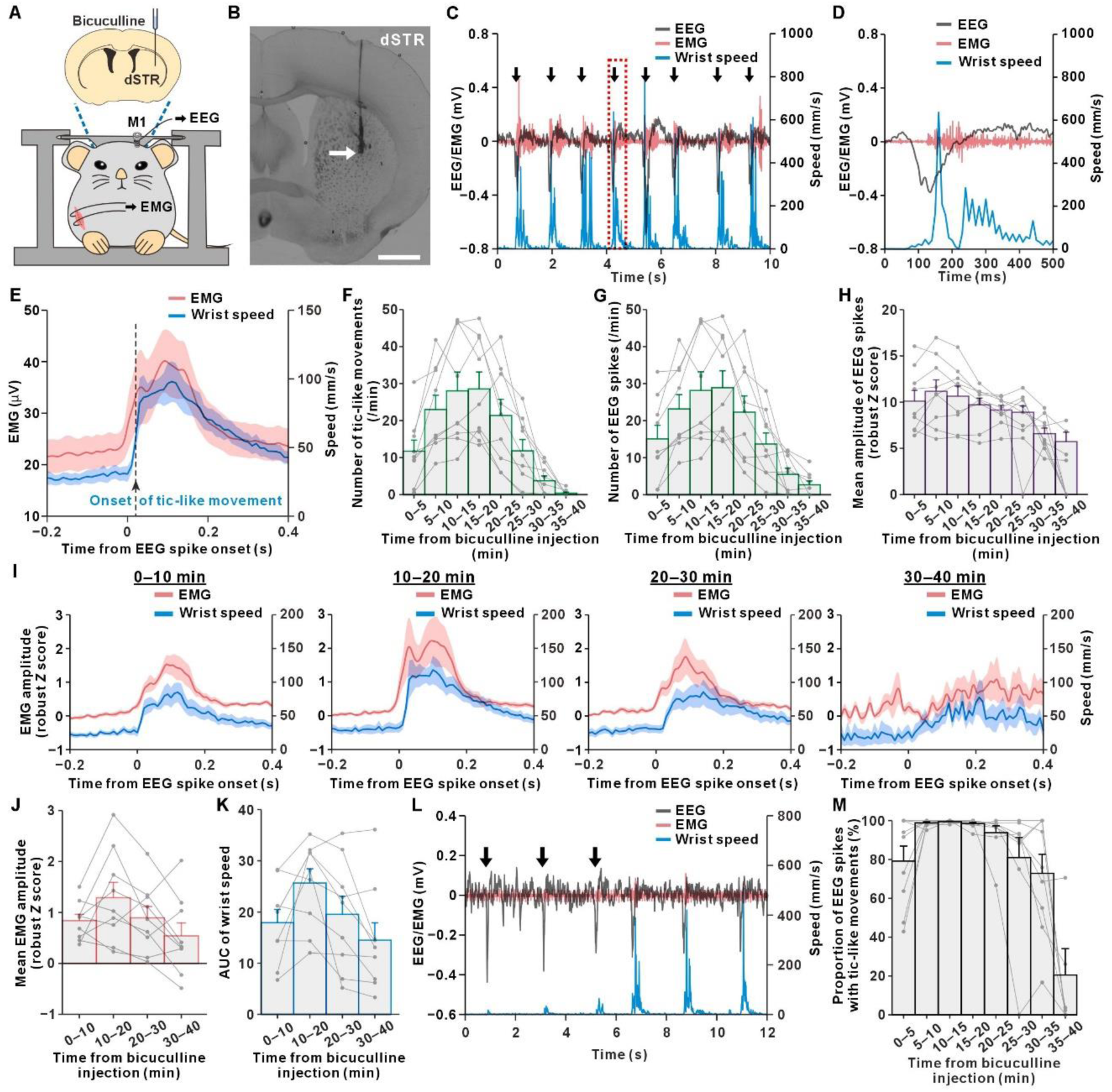
Striatal disinhibition induces tic-like movements and tic-related EEG spikes in the ipsilateral primary motor cortex (M1). (A) Schematic of electrophysiological recordings in the unilateral striatal bicuculline-induced tic mouse model. (B) Coronal brain section showing the injection site in the dorsal striatum (dSTR; arrow). Scale bar, 500 µm. (C and D) Representative traces of electroencephalogram (EEG) from the ipsilateral M1 (gray), electromyogram (EMG) from the contralateral triceps muscle (red), and wrist movement speed of the contralateral forelimb (blue) following bicuculline injection. Black arrows indicate tic-like movement onsets. The region outlined by the red dashed rectangle in (C) is shown at higher magnification in (D). (E) Averaged traces of EMG from the contralateral triceps muscle (pink) and wrist movement speed (blue), aligned to the onsets of EEG spikes recorded in the ipsilateral M1 during the 40-min period following bicuculline injection (n = 9 mice). The black arrow indicates the mean latency of tic onsets relative to EEG spike onsets. Shaded areas represent mean ± SEM. (F-H) Time course of tic counts (F), frequency (G) and mean amplitude (H) of EEG spikes recorded in the ipsilateral M1, plotted in 5-min bins after bicuculline injection (n = 9 mice). (I) Averaged traces of EMG from the contralateral triceps muscle (pink) and wrist movement speed (blue), aligned to the onsets of EEG spikes recorded in the ipsilateral M1, plotted separately for successive 10-min epochs following bicuculline injection (n = 9 mice). Data are presented in the same format as in (E). (J and K) Time course of mean EMG amplitude (J) and the area under the curve (AUC) of the contralateral wrist movement speed (K) within a 0–0.3 s time window following EEG spikes associated with tics, plotted in 10-min bins after bicuculline injection (n = 9 mice). (L) Representative traces showing EEG spikes in the ipsilateral M1 not accompanied by tic-like movements (black arrows). (M) Time course of the percentage of EEG spikes accompanied by tic-like movements, plotted in 10-min bins after bicuculline injection (n = 9 mice). For (H) and (M), if no EEG spikes occurred in a given time bin, the value for that bin was set to 0 for individual data points; however, these bins were excluded from group means and SEMs. Data are presented as mean ± SEM; individual mouse data are shown in gray. See also Figure S1 and Video S1.

We next examined the temporal dynamics of tic-like movements during 40 min following injection. The number of tic events and EEG spikes peaked at 10–20 min, gradually declined after 25 min, and returned to baseline by 40 min (Figures 1F and 1G). The amplitude of EEG spikes remained relatively stable before progressively decreasing (Figure 1H). EMG amplitude and the speed of wrist and mouth movements, quantified in 10-min bins, robustly increased up to 30 min post-injection before markedly attenuating (Figures 1I and S1F). The overall intensity of tic-like movements, estimated from mean EMG amplitude and the area under the curve (AUC) of wrist movement speed (10-min bins), peaked at 10 – 20 min and diminished substantially by 30–40 min (Figures 1J, 1K, and S1G). Non-tic-associated EEG spikes became more prominent after 35 min (Figures 1L and 1M). Thus, striatal disinhibition induced transient tic-like movements lasting ∼40 min in this model.

### Tic-associated brain activation in motor and limbic structures

To identify brain regions engaged during tic expression, we performed c-Fos immunohistochemistry following unilateral striatal bicuculline injection (0.2 µg/µl, 0.20 µl; the same dose as in the preceding experiments). Mice were perfused 90 min after injection, and brain sections were processed for c-Fos staining (Figure 2A). c-Fos mapping revealed robust neuronal activation in multiple brain structures ipsilateral to the injection site (Figures 2A-2C). In the cortex, strong c-Fos expression was observed in medial prefrontal regions (including the cingulate [Cg1], prelimbic [PrL], infralimbic [IL], and dorsal peduncular [DP] cortices), M1, the secondary motor cortex (M2), and IC. Subcortical activation was evident in the BG, including the dSTR, globus pallidus (GP), and subthalamic nucleus (STN). Thalamic activation involved the motor thalamus (ventral anterior [VA] and ventrolateral [VL] nuclei), the sensory thalamus (ventral posterolateral [VPL], ventral posteromedial [VPM], and posterior [Po] nuclei), and the rostral intralaminar thalamic nuclei (rITN; central medial [CM], paracentral [PC], and centrolateral [CL] nuclei). Beyond sensorimotor regions, limbic structures such as the amygdala and hypothalamus also showed high levels of c-Fos expression. Thus, unilateral striatal disinhibition elicited widespread activation across motor and limbic circuits, with strong ipsilateral dominance (Figure 2D).

**Figure 2.**
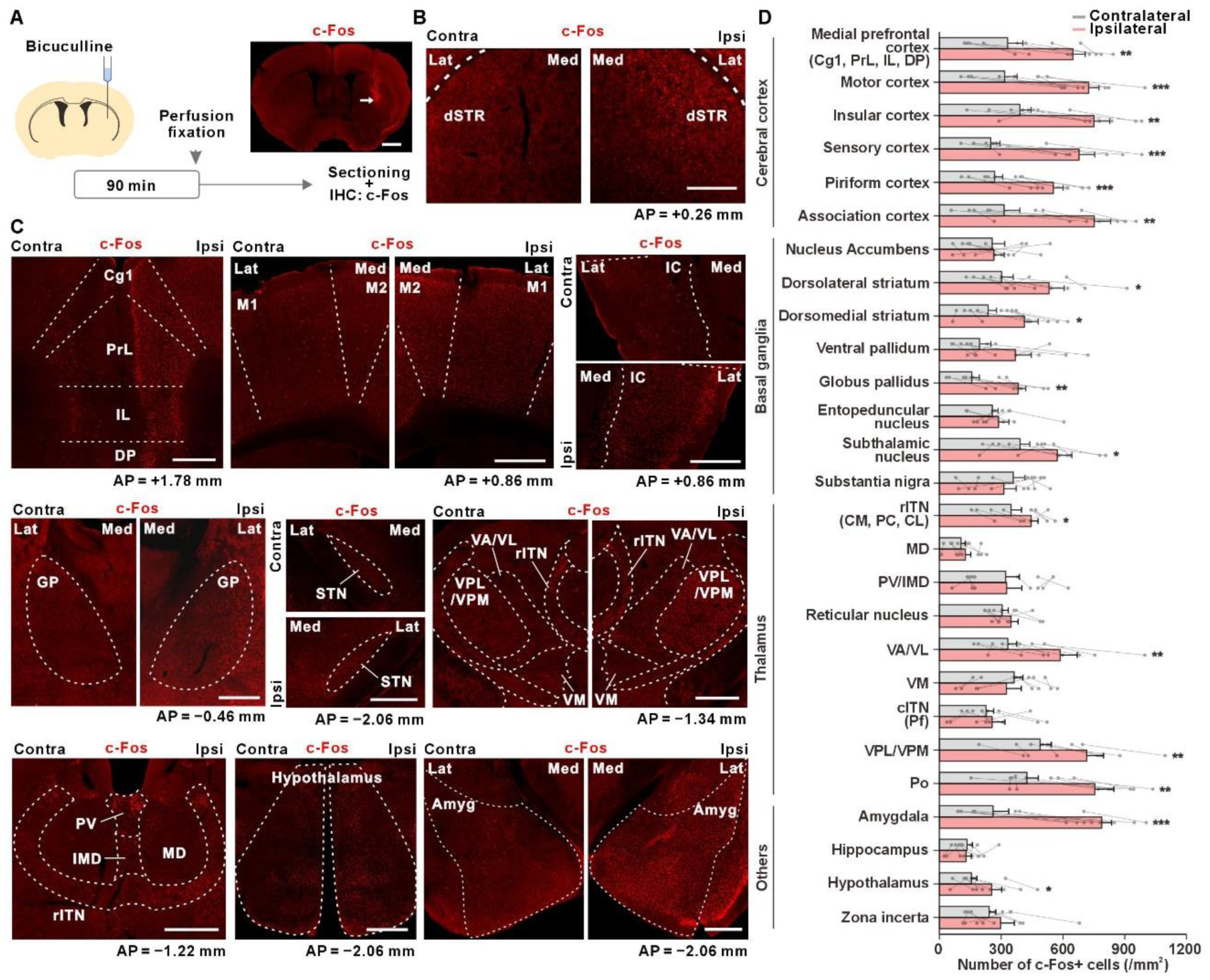
Tic-associated brain activation revealed by c-Fos immunohistochemistry. (A) Experimental timeline. Ninety minutes after unilateral bicuculline injection into the dSTR, mice were perfused, paraformaldehyde-fixed, and brain sections were processed for c-Fos immunohistochemistry. Top right, representative coronal section showing the striatal injection site (arrow) stained for c-Fos. Scale bar, 500 µm. (B) Representative images of c-Fos immunostaining in the contralateral and ipsilateral dSTR at the anteroposterior (AP) level of +0.26 mm from the bregma. Scale bar, 500 µm. (C) Representative c-Fos immunostaining in selected brain regions across cortical, basal ganglia, thalamic, and limbic structures. Coronal sections at the indicated AP levels are shown. Scale bars, 500 µm. (D) Quantification of c-Fos-positive cells in the contralateral (gray) and ipsilateral (pink) hemispheres across brain regions. Data are presented as mean ± SEM; individual mouse data are shown in gray (n = 8 mice). Statistical analyses were performed using paired *t*-tests (*p*-values were not corrected for multiple comparisons). *p < 0.05; **p < 0.01; ***p < 0.001. Abbreviations: Amyg, amygdala; Cg1, cingulate cortex area 1; cITN, caudal intralaminar thalamic nuclei; CL, centrolateral thalamic nucleus; CM, central medial thalamic nucleus; DP, dorsal peduncular cortex; GP, globus pallidus; IC, insular cortex; IL, infralimbic cortex; IMD, intermediodorsal thalamic nucleus; M2, secondary motor cortex; MD, mediodorsal thalamic nucleus; PC, paracentral thalamic nucleus; Pf, parafascicular thalamic nucleus; Pir, piriform cortex; Po, posterior thalamic nuclear group; PrL, prelimbic cortex; PV, paraventricular thalamic nucleus; rITN, rostral intralaminar thalamic nuclei; STN, subthalamic nucleus; VA, ventral anterior thalamic nucleus; VL, ventrolateral thalamic nucleus; VM, ventromedial thalamic nucleus; VPL, ventral posterolateral thalamic nucleus; VPM, ventral posteromedial thalamic nucleus.

### Insular cortical activity is associated with tic generation

Given the widespread activation observed during tic generation, we next focused on IC, since several studies have proposed it as a potential trigger site for tics and their preceding “urges”.^25,26^ Consistent with this, human imaging studies have reported IC activation in TS patients.^27,28^

To monitor tic-related activity in IC and M1 (the final cortical output node for motor commands), we performed *in vivo* Ca^2+^ imaging using fiber photometry (Figures 3A and 3B). Optical fibers were implanted into the ipsilateral M1 and bilateral IC one week after viral injections of AAV1-Syn-GCaMP6f (Figure 3A). Following unilateral striatal disinhibition, robust Ca^2+^ activity was observed in the ipsilateral M1 and IC, time-locked to motor tics (Figures 3B and 3C). The contralateral IC also exhibited tic-associated activity. Analysis of temporal dynamics revealed that tic-associated Ca^2+^ activity in the ipsilateral M1 and bilateral IC peaked at 10–20 min post-injection and then gradually declined (Figures 3D and 3E), paralleling the time course of tic-like movements and M1 EEG spikes (Figures 1F-1K, S1F, and S1G).

**Figure 3.**
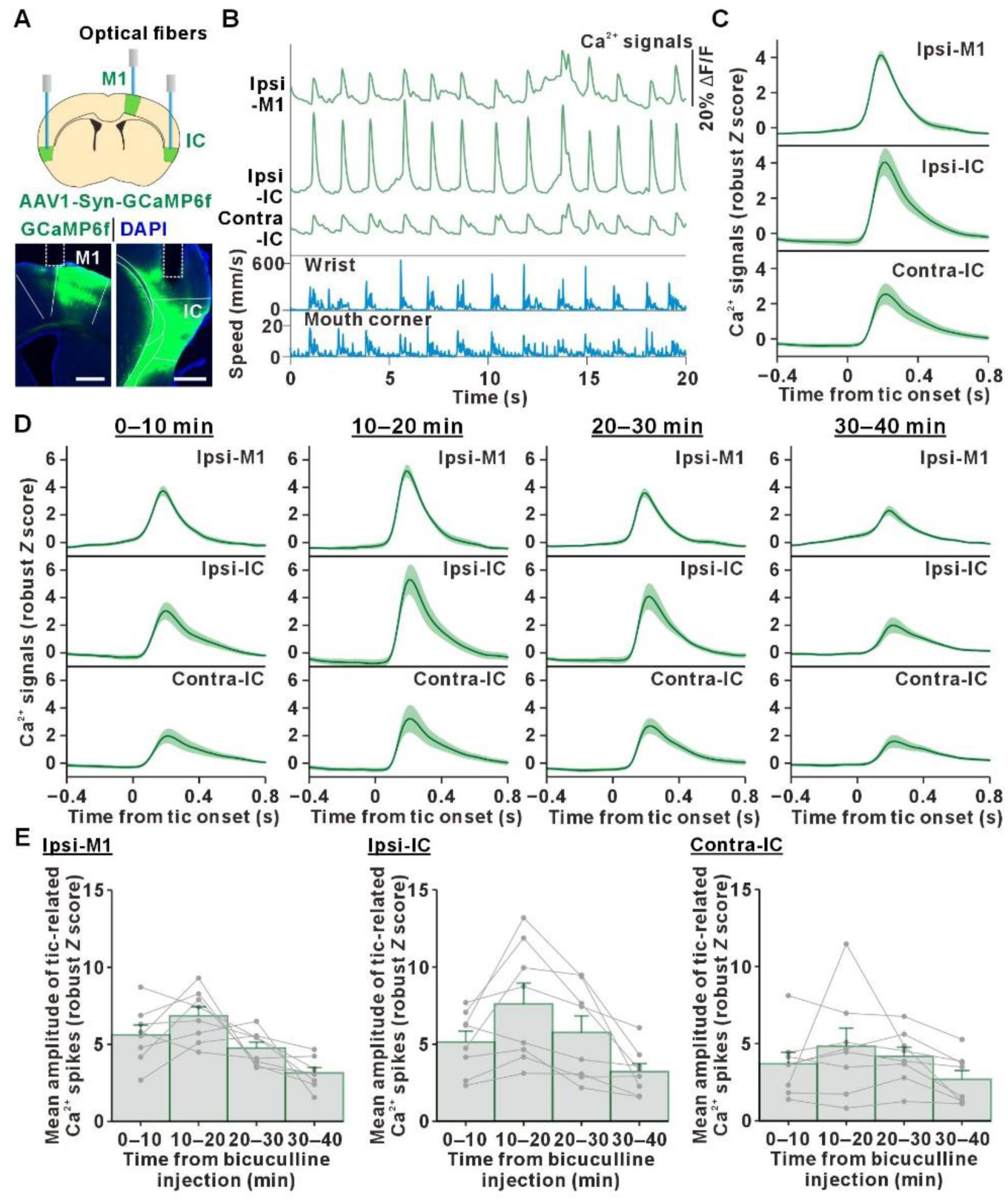
Insular cortical activity is associated with tic generation. (A) Schematic of fiber photometry recordings (top). AAV1-Syn-GCaMP6f was injected into the ipsilateral M1 and bilateral IC. Representative coronal sections showing GCaMP6f-expression and optic fiber placements (dashed lines) in M1 and IC (bottom). Scale bars, 500 µm. (B) Representative traces of Ca^2+^ signals in the ipsilateral M1, ipsilateral IC, and contralateral IC, together with wrist and mouth-corner movement speeds, following unilateral bicuculline injection, (C) Averaged Ca²⁺ signals in the ipsilateral M1, ipsilateral IC, and contralateral IC, aligned to tic onsets during the 40-min post-injection period (n = 8 mice). (D) Same data as in (C), plotted separately for successive 10-min epochs following bicuculline injection. Shaded areas represent mean ± SEM. (E) Time course of mean amplitudes of tic-related Ca²⁺ spikes in the ipsilateral M1, ipsilateral IC, and contralateral IC, plotted in 10-min bins after bicuculline injection. Data are presented as mean ± SEM; individual mouse data are shown in gray.

### The intralaminar thalamic nuclei relay basal ganglia outputs to the insular cortex

We next investigated the neuronal pathway mediating IC activation following striatal disinhibition. To identify outputs from the dSTR, we first injected AAV1-CAG-tdTomato into the dSTR (Figure 4A). Anterogradely labeled axonal terminals were observed in the GP, entopeduncular nucleus (EP), and substantia nigra pars reticulata (SNr), but not in IC or other cortical regions.

**Figure 4.**
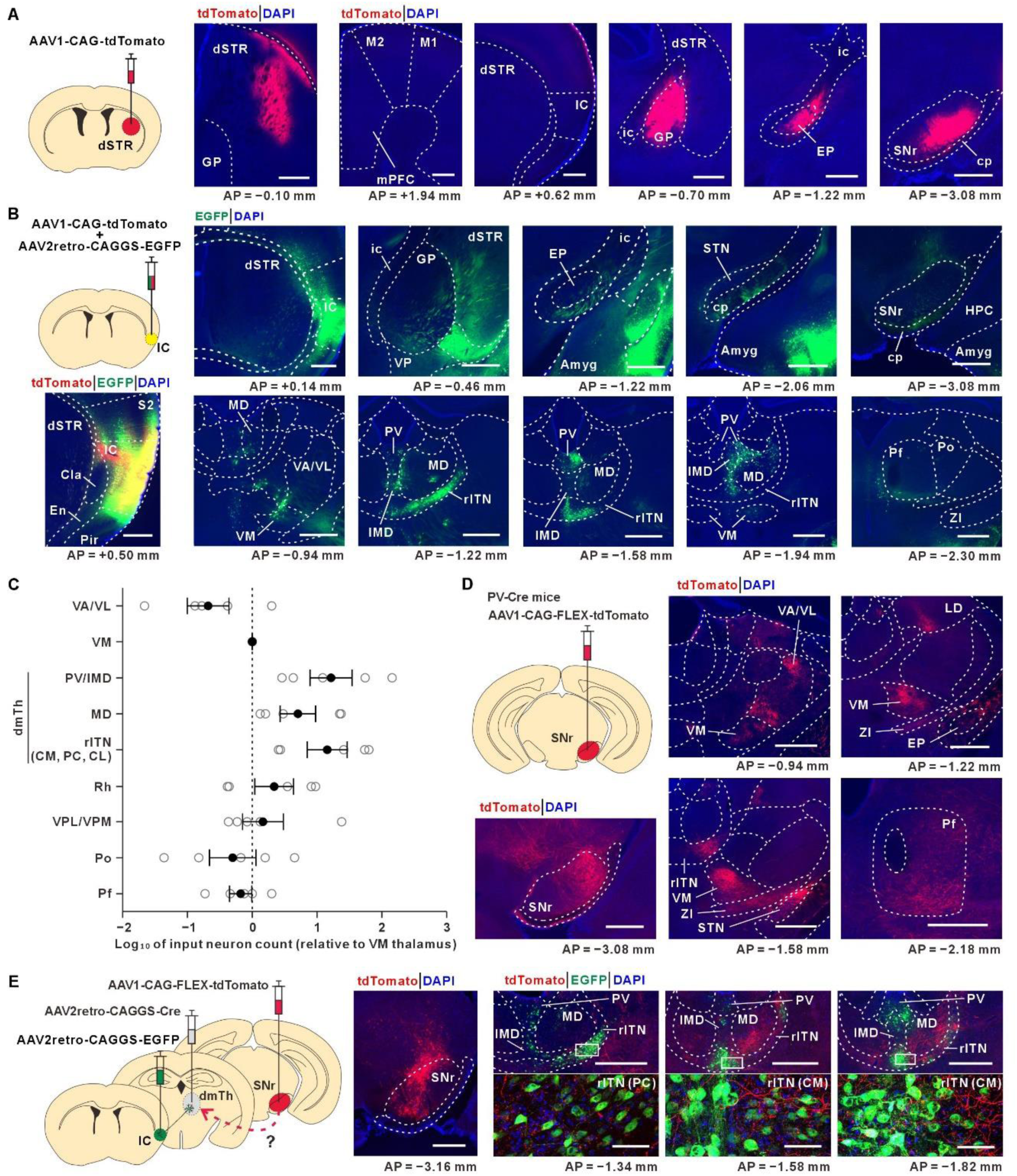
The intralaminar thalamic nuclei relay basal ganglia output to the insular cortex. (A) Anterograde tracing from the dSTR. Injection of AAV1-CAG-tdTomato into the dSTR labeled axon terminals in GP, entopeduncular nucleus (EP), and substantia nigra pars reticulata (SNr), but not in M1, M2, or IC. Scale bars, 500 µm. (B) Retrograde tracing from IC. Injection of AAV2retro-CAGGS-EGFP into the middle IC (along the rostrocaudal axis) labeled neurons in the dorsomedial thalamus (dmTh), including the rITN, PV/intermediodorsal thalamic nuclei (IMD), and MD, but not in the striatum or other basal ganglia nuclei (GP, EP, STN, and SNr). Minimal labeling was observed in the motor thalamus (VM, and VA/VL) and Pf. tdTomato fluorescence in IC (labeled by AAV1-CAG-tdTomato) is shown for reference as the injection site. Scale bars, 500 µm. (C) Quantification of thalamic inputs to IC, expressed as the log_10_ ratio of labeled neurons relative to the number of neurons in VM thalamus (n = 5 mice). Data are presented as mean (black dots) ± SEM. (D) Anterograde tracing from parvalbumin (PV)-positive neurons in SNr of PV-Cre mice. Injection of AAV1-CAG-FLEX-tdTomato revealed widespread projections to multiple thalamic regions, including the dmTh. Scale bars, 500 µm. (E) Triple viral tracing strategy to assess connectivity between SNr and IC via the dmTh. AAV2retro-CAGGS-Cre was injected into the dmTh, AAV1-CAG-FLEX-tdTomato into SNr, and AAV2retro-CAGGS-EGFP into IC. SNr axon terminals (tdTomato, red) were apposed to the somata of IC-projecting dmTh neurons (EGFP, green), particularly within the rITN. White rectangles in the thalamic regions in the top-right panels (scale bars, 500 µm) indicate areas shown at higher magnification in the bottom-right panels (scale bars, 50 µm), highlighting these appositions. Abbreviations: Cla, claustrum; cp, cerebral peduncle; En, endopiriform nucleus; HPC, hippocampus; ic, internal capsule; LD, laterodorsal thalamic nucleus; mPFC, medial prefrontal cortex; Rh, rhomboid thalamic nucleus; S2, secondary somatosensory cortex; VP, ventral pallidum; ZI, zona incerta. See also Figure S2.

To confirm the absence of a direct dSTR-IC projection, we injected AAV2retro-CAGGS-EGFP into the middle part of IC along the rostrocaudal axis (Figure 4B). No retrogradely labeled neurons were found in the striatum or in other BG nuclei (GP, EP, STN, and SNr). Instead, labeled neurons were detected in the dorsomedial thalamus (dmTh), including the rITN, the paraventricular/intermediodorsal nuclei (PV/IMD), and the mediodorsal nucleus (MD) (Figure 4B and 4C). Notably, thalamic structures classically receiving BG outputs, such as the motor thalamus (VA, VL, and ventromedial [VM] nucleus) and the parafascicular nucleus (Pf),^29–31^ contributed minimally to IC-projecting neurons.

Because the SNr is the largest BG output nucleus in rodents,^31^ we hypothesized that SNr neurons reach IC via the dmTh. A large proportion of SNr neurons are PV-positive and exhibit projections similar to VGAT-positive neurons,^31,32^ indicating that they are GABAergic projection neurons. We injected AAV1-CAG-FLEX-tdTomato into SNr of PV-Cre mice (Figure 4D), revealing widespread projections to multiple thalamic regions, including the dmTh (Figures 4D and S2).

To test whether dmTh-projecting SNr neurons connect with IC-projecting dmTh neurons, we performed triple viral injections: 1) AAV2retro-CAGGS-Cre in the dmTh, 2) AAV1-CAG-FLEX-tdTomato into SNr, and 3) AAV2retro-CAGGS-EGFP into IC (Figure 4E). SNr axon terminals were closely apposed to IC-projecting dmTh neurons, particularly within the rITN.

These findings indicate that IC receives BG outputs via ITN, known targets of deep brain stimulation (DBS) in TS patients.^33,34^

### Chemogenetic inhibition of thalamo-insular pathway alleviates tic-like movements

To determine whether the thalamo-insular pathway contributes to motor tic generation, we tested its role using designer receptors exclusively activated by designer drugs (DREADD). We then examined whether tic-like movements induced by striatal disinhibition were attenuated.

As an initial step, we examined whether direct inhibition of IC reduced tic expression. AAV8-hSyn-hM4Di-mCherry was bilaterally injected into three rostrocaudally distinct IC sites (Figures 5A and 5C). Thirty minutes after intraperitoneal administration of either vehicle or clozapine-N-oxide (CNO; DREADD agonist), bicuculline was injected into the same dSTR site via a guide cannula across three test sessions. To control for repeated injection effects, the sequence was vehicle (1st test), CNO (2nd test), and vehicle (3rd test) (Figure 5E). Tic severity was quantified during the 30-min period following bicuculline injection, when tics were prominent, by measuring both the number of tics and the intensity of tic-associated body movements, defined by movement speed (see STAR Methods). Chemogenetic inhibition of bilateral IC significantly reduced both tic frequency and intensity (Figures 5F and 5G; Video S2).

**Figure 5.**
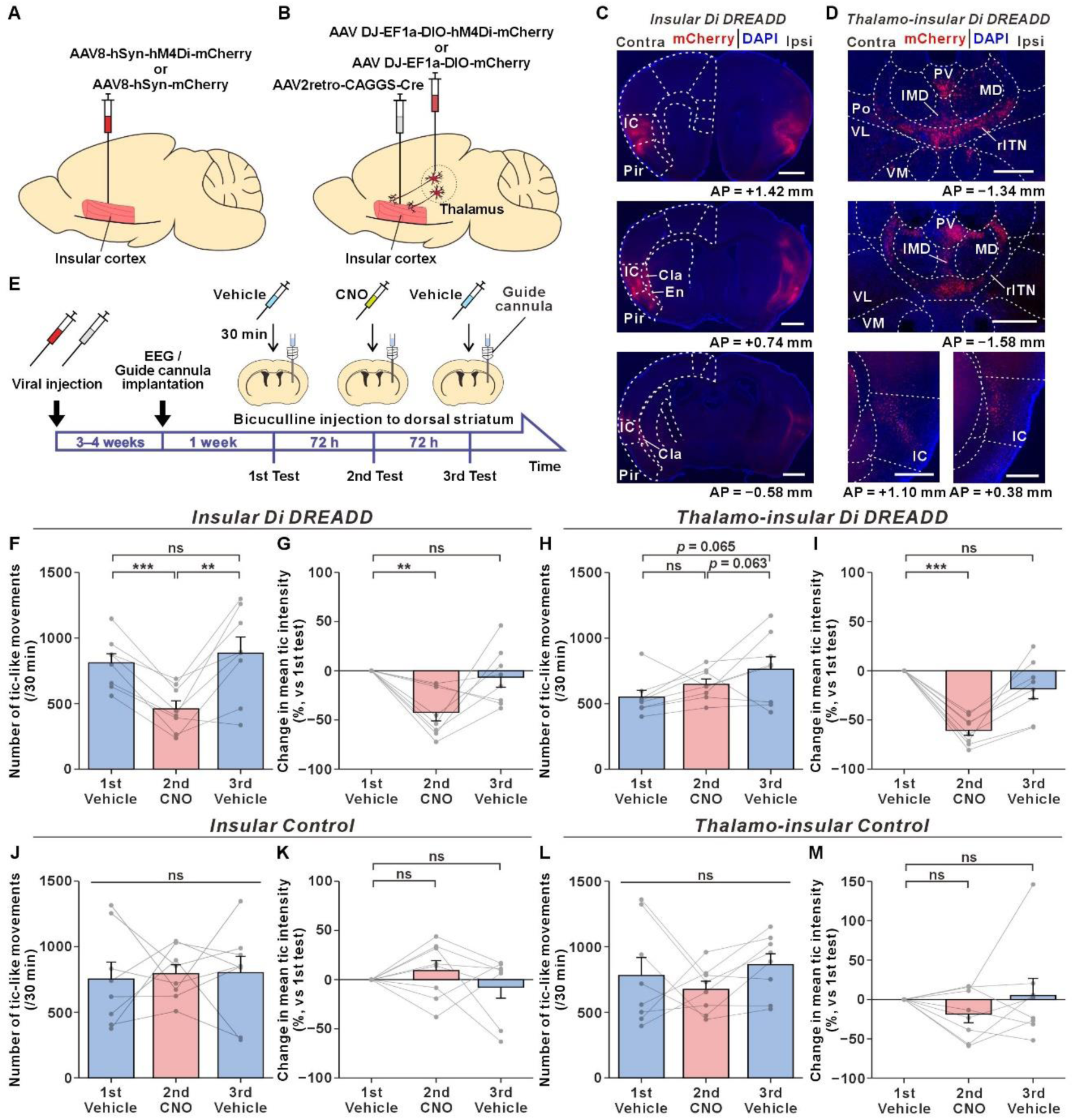
Chemogenetic inhibition of the thalamo-insular pathway alleviates tic-like movements. (A and B) Schematics of the experimental setup for chemogenetic inhibition of IC or the dmTh-IC pathway. To inhibit IC, AAV8-hSyn-hM4Di-mCherry (or control AAV8-hSyn-mCherry) was bilaterally injected into three rostrocaudally distinct IC sites. To specifically inhibit the thalamo-insular pathway, AAV2retro-CAGGS-Cre was bilaterally injected into IC and AAV-DJ-EF1α-DIO-hM4Di-mCherry (or control AAV-DJ-EF1α-DIO-mCherry) into bilateral dmTh. (C and D) Representative coronal sections showing hM4Di-mCherry-expression in IC (C) or in dmTh and IC (D). Scale bars, 500 µm. (E) Experimental timeline. Mice received repeated unilateral bicuculline injections into the dSTR via a chronically implanted guide cannula, following intraperitoneal administration of vehicle (1st test), clozapine-N-oxide (CNO, 10 mg/kg; 2nd test), and vehicle (3rd test). (F and G) Chemogenetic inhibition of bilateral IC reduced both tic frequency (F) and tic intensity (G) during the 30-min period after bicuculline injection (n = 8 mice). (H and I) Chemogenetic inhibition of bilateral dmTh-IC pathways reduced tic intensity (I), but not tic frequency (H) (n = 8 mice). (J-M) Control experiment using mCherry-expressing viruses in bilateral IC (J, K) or bilateral dmTh-IC pathways (L, M) showed no effects of CNO administration. (n = 8 mice for each group). (F-M) Data are presented as mean ± SEM; individual mouse data are shown as gray circles. Statistical analyses were performed using one-way repeated measures ANOVA followed by pairwise *t*-tests with Holm correction (F, H, J, and L) or one-sample *t*-tests with Holm correction (G, I, K, and M). ns, not significant; **p < 0.01; ***p < 0.001. See also Video S2.

We next investigated whether chemogenetic inhibition of the dmTh-IC pathway affected tic generation. To address this issue, AAV2retro-CAGGS-Cre was injected into bilateral IC and AAV-DJ-EF1α-DIO-hM4Di-mCherry into bilateral dmTh (Figures 5B and 5D). Bilateral inhibition of this pathway significantly reduced tic intensity, while tic frequency was unchanged (Figures 5H and 5I; Video S2).

Because we used a relatively high dose of CNO (10 mg/kg) and CNO has been reported to influence glutamatergic and dopaminergic systems,^35,36^ we conducted control experiments using mCherry-expressing viruses instead of hM4Di-mCherry (Figures 5A and 5B). CNO administration in these controls had no significant effect (Figures 5J-5M).

Taken together, these results indicate that the thalamo-insular pathway contributes to tic generation.

### Chemogenetic inhibition of the thalamo-insular pathway alters downstream tic-associated cortical activity

We next examined whether chemogenetic manipulation of the thalamo-insular pathway altered activity in downstream cortical regions. Fiber photometry Ca^2+^ recordings were performed in the ipsilateral M1 and bilateral IC of mice in which IC-projecting dmTh neurons expressed inhibitory DREADD, using the same experimental timeline as in the preceding experiment (Figure 6A). CNO administration reduced tic intensity without affecting tic frequency, consistent with behavioral data (Figures 5H, 5I, and S3). During the CNO session, tic-associated Ca^2+^ activity persisted (Figures 6B and 6C), but its amplitude was significantly reduced in the ipsilateral M1 and ipsilateral IC (Figures 6C and 6D). In contrast, the amplitude in the contralateral IC remained unchanged.

**Figure 6.**
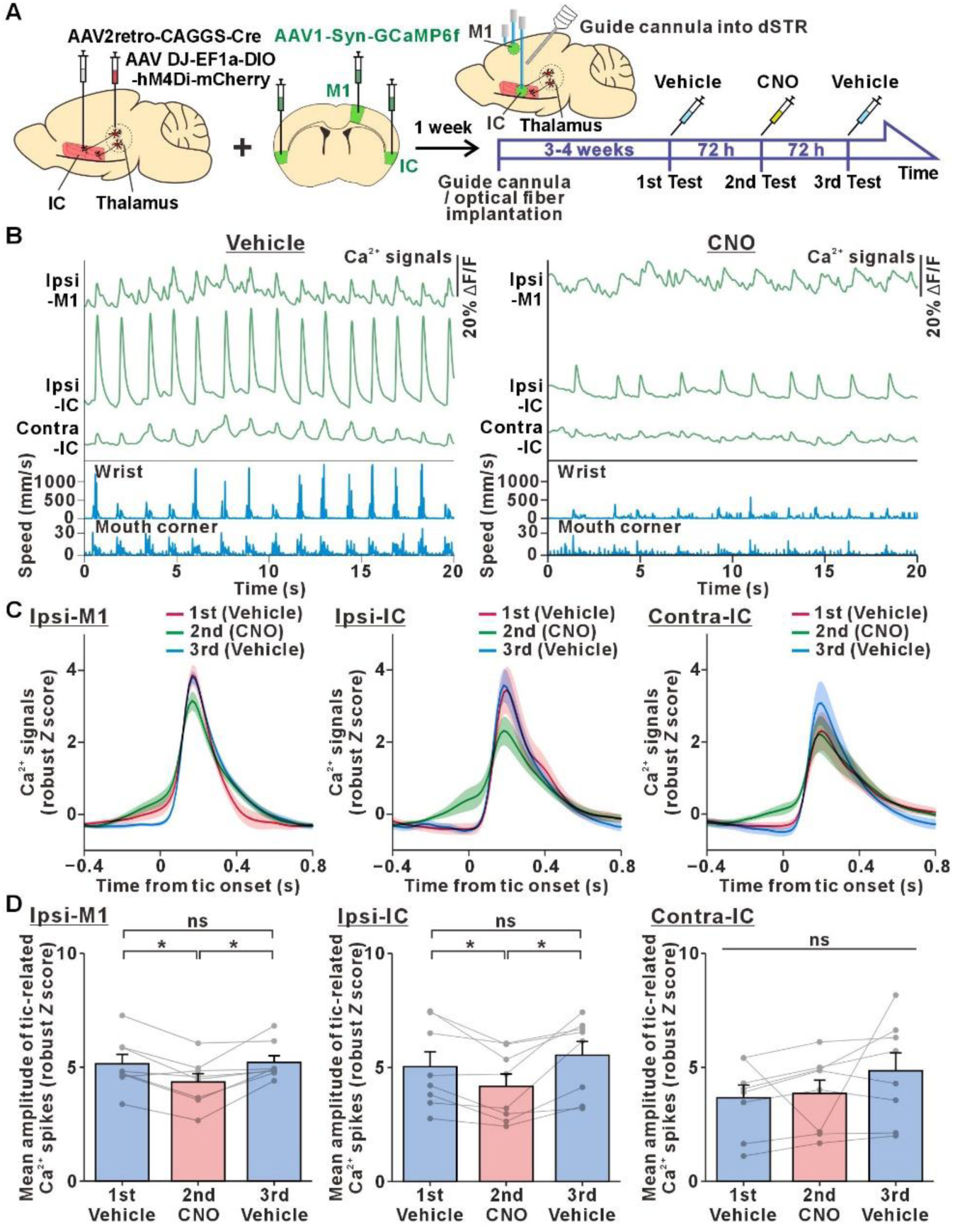
Chemogenetic inhibition of the thalamo-insular pathway alters downstream tic-associated cortical activity. (A) Shematic of the experimental setup for fiber photometry recording during chemogenetic inhibition of the dmTh-IC pathway. To inhibit this pathway, AAV2retro-CAGGS-Cre was bilaterally injected into IC, and AAV-DJ-EF1α-DIO-hM4Di-mCherry (or control AAV-DJ-EF1α-DIO-mCherry) into bilateral dmTh. AAV1-Syn-GCaMP6f was injected into the ipsilateral M1 and bilateral IC for fiber photometry recordings. Following implantation of a guide cannula and optical fibers, tic-like movements were induced by repeated unilateral bicuculline injections into the dSTR under vehicle (1st test), CNO (10 mg/kg; 2nd test), and vehicle (3rd test) conditions. (B) Representative traces of Ca^2+^ signals in the ipsilateral M1, ipsilateral IC, and contralateral IC, together with wrist and mouth-corner movement speeds, following unilateral bicuculline injection during vehicle (left) and CNO (right) sessions. (C) Averaged tic-aligned Ca^2+^ signals in the ipsilateral M1, ipsilateral IC, and contralateral IC, across the three sessions (n = 8 mice). Shaded areas represent mean ± SEM. (D) Quantification of tic-related Ca^2+^ amplitudes in the ipsilateral M1, ipsilateral IC, and the contralateral IC (n = 8 mice). Chemogenetic inhibition of the dmTh-IC pathway significantly reduced Ca²⁺ amplitudes in the ipsilateral M1 and ipsilateral IC, but not in the contralateral IC. Data are presented as mean ± SEM; individual mouse data are shown as gray circles. Statistical analyses were performed using one-way repeated measures ANOVA followed by pairwise *t*-tests with Holm correction. ns, not significant; *p < 0.05. See also Figures S3-S5.

To further assess whether chemogenetic manipulation of IC or the dmTh-IC pathway modulates tic-associated M1 activity at the single-cell level, we performed two-photon Ca^2+^ imaging of GCaMP6f-expressing ipsilateral M1 neurons (Figures S4A and S4B). Following bicuculline injection, M1 neurons exhibited temporally synchronized Ca^2+^ signals (Figures S4B and S4C; see also Video S3). IC inhibition reduced Ca^2+^ event amplitudes in both superficial (layer 2/3) and deep (layer 5/6) layers, whereas dmTh-IC pathway inhibition selectively reduced amplitudes in the superficial layer. Event frequency in the superficial layer was unaffected by either manipulation, whereas IC inhibition increased frequency in the deep layer. For response probability of M1 Ca^2+^ signals to tic onsets (see STAR Methods), decreases were observed in the superficial layer under both manipulations, whereas dmTh–IC pathway inhibition uniquely led to an increase in the deep layer (Figures S4D-I).

These results indicate that inhibition of IC neurons or IC-projecting dmTh neurons modulates M1 activity in a layer-specific manner, contributing to tic suppression.

## DISCUSSION

We found that a striatal disinhibition-induced mouse model of tics exhibited brain activation across motor and limbic structures, including IC. We further provided anatomical evidence that BG outputs originating from the striatal motor domain are relayed to IC via ITN, which are known targets of DBS in intractable TS.^33,34^ Finally, we demonstrated that chemogenetic inhibition of the thalamo-insular pathway suppressed tic symptoms. Collectively, these findings reveal an unexpected crosstalk between motor and limbic domains within the CBGTC network and suggest that aberrant activity in the thalamo-insular pathway contributes to the emergence of tics.

### Animal model of tic disorders based on striatal disinhibition

Motor and vocal tics accompanied by premonitory urges are hallmark symptoms of TS patients. These individuals frequently manifest comorbid psychiatric and behavioral disorders, including OCD, ADHD, autism spectrum disorder (ASD), anxiety, depression, and sleep disorders.^7,8^ Although recent genome-wide association studies (GWAS) and whole-exome sequencing studies have suggested some candidate genes contributing to TS pathophysiology, critical genes have not yet been identified.^37–39^ Moreover, knockout mouse models of these candidate genes generally do not exhibit motor tics, or, if they do, the frequency is very low. Instead, some of these models display stereotyped behaviors such as excessive grooming or rearing, which more closely resemble OCD-like behaviors.^40,41^ In contrast, postmortem studies in TS patients have revealed a reduction of striatal GABAergic interneurons.^21,22^ Consistent with these findings, nonhuman primate models of motor and vocal tics induced by blocking GABAergic inputs have been well established (see also Figures 1 and S1 in our study).^15–19,42^

In addition to striatal interneurons, the striatum is also known to receive substantial GABAergic inputs from “arkypallidal” neurons in GP (corresponding to the external globus pallidus [GPe] in primates; see also Figure S1A).^43^ Indeed, injections of a GABA_A_ receptor antagonist into primate GPe induce choreic dyskinesias.^44–46^ Thus, dysfunction of the striato-GP network through disruption of GABAergic tone may generate synchronized activity in BG output neurons and subsequently M1 neurons, ultimately leading to involuntary movements.^42,47,48^

### The insula is a key site responsible for tic generation

Our study demonstrated that IC is robustly activated in a mouse model of tics induced by striatal disinhibition (Figure 2). Tic-associated activity was observed in IC as well as in M1 (Figures 3, 6, and S4). Importantly, chemogenetic inhibition of IC alleviated tic-like behaviors (Figure 5). These findings strongly suggest that IC serves as a key site responsible for tic generation. Supporting this notion, human imaging studies in TS patients have demonstrated IC activation.^27,28^

Intriguingly, we previously reported that the use of an oral splint, typically applied in the treatment of temporomandibular disorders, produced immediate and marked therapeutic effects on both motor and vocal tics in approximately two-thirds of TS patients.^49^ Motivated by this clinical observation, we sought to anatomically trace the ascending pathway of orofacial proprioception in rodents, given that oral devices activate muscle spindles of the masseter. We found that orofacial proprioceptive inputs are ultimately transmitted not to the primary sensory cortex but rather to IC, via the supratrigeminal nucleus (Su5) and the caudo-ventromedial subdivision of VPM (VPMcvm).^50,51^ Taken together, these anatomical and clinical findings suggest that oral splint insertion may modulate aberrant IC activity underlying tic generation in TS patients.

IC is well known as a center of emotional processing and interoceptive awareness, integrating signals generated by physiological body states.^52–54^ In this context, abnormal IC activity in TS patients may contribute not only to motor symptoms such as tics but also to premonitory urges and comorbid psychiatric disorders, including OCD, ADHD, and ASD.

### Why the intralaminar thalamic nuclei can be surgical targets in TS patients

Historically, ablative procedures such as thalamotomy, pallidotomy, and subthalamotomy were pioneering neurosurgical treatments for movement disorders.^55–57^ Since the development of DBS at the end of the last century, DBS targeting the internal globus pallidus (GPi) or STN has become the gold standard for intractable Parkinson’s disease and dystonia.^58–60^ In contrast, ITN have been empirically identified as effective DBS targets in TS patients.^61,62^ However, the mechanistic basis for this efficacy has remained unclear.

Anatomically, our study identified the dmTh, particularly the rITN, as a candidate relay region of BG outputs to IC (Figure 4). Chemogenetic inhibition of IC-projecting dmTh neurons attenuated tic-associated activity in IC (Figure 6). Moreover, this manipulation also suppressed tic-related activity in M1 and reduced tic-like movements (Figures 5, 6, S3, and S4). These results suggest that the dmTh-IC pathway transmits abnormal BG activity to IC, thereby contributing to the exacerbation of motor tics.

Importantly, the caudal ITN (cITN), a clinical DBS target in TS patients, maintains topographically organized connections with the CBGTC circuits (particularly the striatum and cortex).^30,63^ In contrast, the rITN exhibits widespread or loosely topographical projections to multiple structures within these circuits.^64^ Given that the rITN and the cITN are spatially adjacent, the current spread from a DBS electrode implanted in the cITN may also influence the rITN, thereby exerting therapeutic effects in TS patients. A future double-blinded clinical trial directly comparing the rITN versus cITN stimulation will be warranted to determine the optimal surgical target.

### Crosstalk in motor and limbic networks in the cortico-basal ganglia-thalamo-cortical circuits

The existence of crosstalk between motor and limbic networks within the CBGTC circuits has long been debated. These circuits have traditionally been conceived as functionally segregated, closed parallel loops, comprising motor, associative, and limbic loops that originate from distinct cortical regions.^65,66^ Consistent with this view, pharmacological manipulations in the motor, associative, and limbic domains of the striatum or GPe selectively induce motor, cognitive, and emotional disturbances, respectively.^19,45,67^ Although IC is not included in the classical definitions of the limbic system proposed by Papez,^68^ converging evidence places IC within a broader limbic network that links emotion, interoception, and behavior.^52,69^ Importantly, IC directly projects to the ventral striatum (striatal limbic domain).^70,71^

Our findings challenge this closed-loop model. In addition to the previously described “limbic-to-motor” open loop (striatal limbic domain-SNr-motor thalamus-motor cortices pathway),^32^ we identified a complementary “motor-to-limbic” open loop (striatal motor domain-SNr-dmTh-IC pathway; see also Figure S5). These subcortical interactions indicate that motor and limbic domains within the CBGTC circuits are not strictly segregated, but instead communicate bidirectionally. While the closed “motor cortices-BG motor domain-motor thalamus-motor cortices” loop may account for the pathophysiology of simple motor tics, our new framework more effectively explains the broad spectrum of symptoms observed in TS patients, including motor and vocal tics, premonitory urges, and psychiatric comorbidities such as OCD and ADHD. Dysfunction in the “motor-to-limbic” pathway may permit abnormal motor signals to spread into limbic regions, leading to emotional and cognitive symptoms. In this context, targeted intervention of aberrant neuronal processing in the “motor-to-limbic” circuit (i.e., the dmTh-IC pathway) could be therapeutically beneficial. Approaches such as ultrasound neuromodulation or thermoablation, which can access deep brain structures, may provide therapeutic benefits in TS patients. In contrast, interventions such as transcranial direct current or transcranial magnetic stimulation, when applied to M1, may inadvertently disrupt voluntary motor control.^72–74^

Beyond subcortical interactions, motor-limbic crosstalk may also occur at the cortical level. The limbic cortex (e.g., IC) has limited direct connections to M1,^71^ but aberrant IC activity may influence motor outputs indirectly through other cortical structures, such as the sensory cortex, or through subcortical feedback loops.^32^ Thus, motor and limbic loops communicate at both subcortical and cortical levels, and abnormal dynamics across these multiple pathways may underlie the recurrent and dynamic nature of tic generation.

In conclusion, our study suggests that normalization of IC activity could restore balance between motor and limbic circuits and improve both motor and emotional symptoms in TS.

## RESOURCE AVAILABILITY

### Lead contact

Further information and requests for resources and reagents should be directed to the lead contact, Yoshihisa Tachibana.

### Materials availability

The study did not produce new unique reagents.

### Data and code availability

Tic-associated raw data have been deposited at Mendeley Data. Accession numbers are listed in the key resources table. This paper does not report original code. Any additional information required to reanalyze the data reported in this paper is available from the lead contact upon request.

## ACKNOWLEDGMENTS

We thank A. Nambu, S. Yamamoto, K. Toda, T. Hirata, K. Tamada, N. Nakai, and Y. Shintani for their valuable comments and discussions. We are grateful to H. Shima and M. Shiramizu for technical assistance with animal care. This work was supported by the Japan Society for the Promotion of Science (JSPS) KAKENHI (18K06852, 22K19732, 24H00422, and 24K02339 to Y.T.; 24H00620 to T.T.), by the Taiju Life Social Welfare Foundation (to Y.T.), and by AMED under Grant Number JP23wm0625001 (to K.K.).

## AUTHOR CONTRIBUTIONS

H.K. and Y.T. conceived the study and designed the experiments. H.K. performed the majority of the experiments, and N.T. conducted some of the histological experiments. H.K. analyzed the data. K.K. provided materials. H.K. and Y.T. wrote the manuscript with editorial input from T.T.

## DECLARATION OF INTERESTS

The authors declare no competing interests.

## EXPERIMENTAL MODEL AND SUBJECT DETAILS

### Animals

All animal procedures were approved by the Institutional Animal Care and Use Committees of Kobe University. Male C57BL/6J (6–16 weeks old) and male PV-Cre mice^75^ (6–10 weeks old) were used. All animals were maintained in the animal facility at Kobe University. Animals were group-housed under controlled temperature (23–25℃) in a 12/12 h light-dark cycle (lights on from 6 AM to 6 PM) with free access to food and water. Animals implanted with a guide cannula or optical fibers were single-housed after surgery to prevent damage to the implanted devices.

## METHOD DETAILS

### Head plate attachment

The first surgery was performed on mice aged 6–10 weeks. Mice were anesthetized with ketamine (80 mg/kg, intraperitoneally [i.p.]; Ketalar, Daiichi Sankyo, Tokyo, Japan) and xylazine (12 mg/kg, i.p.; Seractar, Elanco Japan, Tokyo, Japan), and mounted on a stereotaxic apparatus (SR-6M-HT, Narishige, Tokyo, Japan). To prevent corneal drying, gentamicin eye ointment (TAKATA Pharmaceutical Co., Saitama, Japan) was applied to both eyes. After subcutaneous injection of epinephrine-containing lidocaine (Xylocaine DENTAL, Dentsply Sirona, York, PA, USA), the scalp was incised to expose the skull. Except for anatomical neuronal tracing experiments, a custom-made metal head plate was affixed to the skull with dental cement (G-CEM ONE neo, GC.dental, Tokyo, Japan) to secure the animal in a custom-built fixation apparatus.

### Viral injection

For viral delivery, a small craniotomy was performed on the same day as head plate attachment. A small burr hole was made using a dental drill (MICROMOTOR LM-III, GC.dental), with supplementary ketamine/xylazine administered as needed. Viral solutions were pressure-injected through a glass micropipette attached to a microinjector (I-31, Narishige, Tokyo, Japan). The micropipette was left in place for at least 5 min to minimize backflow and prevent viral spread to adjacent brain regions. All stereotaxic coordinates were determined according to the Paxinos Mouse Brain Atlas (third edition).^76^

For viral tracing experiments, AAV1-CAG-tdTomato (2.0 × 10^13^ vector genomes [vg]/ml, diluted 1:2 in saline) or AAV1-CAG-Flex-tdTomato (1.6 × 10^13^ vg/ml, diluted 1:2 in saline) was injected unilaterally into the dorsal striatum (dSTR; 0.10–0.15 µl/injection, AP: 0.0 mm, ML: 2.7 mm, DV: –3.5 mm from the bregma) or the substantia nigra pars reticulata (SNr; 0.10–0.15 µl/injection, AP: –3.1 mm, ML: 1.65 mm, DV: –4.6 mm). AAV2retro-CAGGS-EGFP (6.8 × 10^12^ vg/ml, diluted 1:2 in saline) or a mixture of AAV2retro-CAGGS-EGFP and AAV1-CAG-tdTomato (diluted 1:1:1 in saline; red fluorescence data not shown) was injected into the middle portion of the insular cortex (IC) along the rostrocaudal axis (0.10–0.15 µl/injection; AP: 0.5 mm, ML: 3.3 mm, DV: –3.6 mm). For labeling dorsomedial thalamus (dmTh)-projecting neurons in other brain regions, AAV2retro-CAGGS-Cre (7.3 × 10^12^ vg/ml, diluted 1:2 in saline) was injected into the dmTh (0.20 µl/injection, AP –1.5 mm, ML: 0.5 mm, DV: –3.5 mm).

For experiments targeting IC neurons with hM4Di-mCherry or mCherry using the designer receptors exclusively activated by designer drugs (DREADD) system, AAV8-hSyn-hM4Di-mCherry (2.2 × 10^13^ vg/ml, diluted 1:2 in saline) or AAV8-hSyn-mCherry (2.1 × 10^13^ vg/ml, diluted 1:2 in saline) was injected bilaterally into three rostrocaudally distinct IC sites: anterior IC (aIC; 0.20 µl/injection, AP: 1.5 mm, ML: ±2.7 mm, DV: –3.2 mm), middle IC (mIC; 0.30 µl/injection, AP: 0.5 mm, ML: ±3.3 mm, DV: –3.6 mm), and posterior IC (pIC; 0.20 µl/injection, AP: –0.5 mm, ML: ±3.7 mm, DV: –3.5 mm).

For experiments targeting IC-projecting dmTh neurons with hM4Di-mCherry or mCherry, AAV2retro-CAGGS-Cre (7.3 × 10^12^ vg/ml, diluted 1:2 in saline) was injected bilaterally into three distinct IC sites (aIC, mIC, and pIC; same coordinates as above), and AAV-DJ-EF1α-DIO-hM4Di-mCherry (1.5 × 10^13^ vg/ml, diluted 1:2 in saline) or AAV-DJ-EF1α-DIO-mCherry (2.0 × 10^13^ vg/ml, diluted 1:2 in saline) was injected bilaterally into the dmTh (0.40 µl/injection, AP: –1.5 mm, ML: ±0.5 mm, DV: –3.5 mm).

For fiber photometry, AAV1-Syn-GCaMP6f-WPRE-SV40 (2.0 × 10^13^ vg/ml, diluted 1:2 in saline) was injected unilaterally into the primary motor cortex (M1; 0.20 µl/injection, AP: 0.5 mm, ML: 1.0 mm, DV: –1.3 mm) or bilaterally into the mIC (0.20 µl/injection, AP: 0.5 mm, ML: 3.3 mm, DV: –3.6 mm). One week later, optical fibers (1.25 mm ferrule OD, 400 μm core, NA 0.5; cat# R-FOC-BL400C-50NA, RWD Life Science, Guangdong, China) were implanted 200 µm above the injection sites (M1 or mIC) under isoflurane anesthesia (1.0%; Viatris, Pittsburgh, PA, USA) and fixed with dental resin (UNIFAST II, GC.dental).

For two-photon microscopic imaging, a circular craniotomy (2.5 mm in diameter) was made over M1 (center, AP: 0.5 mm, ML: 1.0 mm) ipsilateral to the striatal bicuculline injection (see below) under isoflurane anesthesia, 1-2 days after the head plate attachment. After the craniotomy, AAV1-Syn-GCaMP6f-WPRE-SV40 (2.0 × 10^13^ vg/ml, diluted 1:2 in saline) was injected into two depths of the ipsilateral M1 (300 µm and 500 µm below the brain surface, 0.20 µl at each site). The cortical surface was covered with a glass window composed of two coverslips (2 and 4.5 mm in diameter, respectively; Matsunami Glass, Osaka, Japan), using transparent adhesive (NOR-61, Norland Products, Jamesburg, NJ, USA). The edges of the glass window were sealed with dental cement (Fuji Lute, GC.dental) and adhesive resin cement (Super Bond, Sun Medical, Shiga, Japan).

### EEG and EMG electrode implantation

In the experiments shown in Figures 1 and S1, electroencephalogram (EEG) and electromyogram (EMG) electrodes were implanted under ketamine/xylazine anesthesia on the same day as head plate attachment. In the experiment shown in Figure 5, EEG electrodes were implanted under isoflurane anesthesia, three weeks after viral injections.

For EEG electrode implantation, small burr holes were made above the ipsilateral M1 (AP: 0.5 mm, ML: 1.0 mm) and the contralateral visual cortex (AP: –3.5 mm, ML: 3.0 mm; reference electrode). Metal screws serving as electrodes were implanted into the holes and secured to the skull with dental cement (G-CEM ONE neo, GC.dental). Screws were connected to terminal connectors via copper wires, and the connectors were mounted on the head plate using dental resin (UNIFAST II, GC.dental).

For EMG electrode implantation, mice received a subcutaneous injection of epinephrine-containing lidocaine in the contralateral upper arm. The skin was incised to expose the triceps muscle, and two PFA-coated stainless-steel wires (7-strand, 0.0055” in diameter; cat# 793200, A-M Systems, Sequim, WA, USA) were inserted into the muscle and secured with tissue adhesive (Vetbond, 3M Co., Ltd, St. Paul, MN, USA) and suture thread. The implanted wires were connected to terminal connectors as described above.

### Guide cannula implantation

When repeated bicuculline injections were required for DREADD experiments (see below for details), a guide cannula was chronically implanted into the unilateral dSTR to minimize variability in behavioral outcomes caused by differences in bicuculline injection sites. After a craniotomy under isoflurane anesthesia, a 5-mm-long guide cannula (AG-5T, Eicom, Kyoto, Japan) was implanted vertically into the dSTR (AP: 0.0 mm, ML: 2.7 mm, DV: –3.5 mm) for DREADD experiments not involving neuronal recording. For DREADD experiments combined with fiber photometry or two-photon calcium imaging, an 8-mm-long guide cannula (CXG-8T, Eicom) was implanted sagittally from caudal to rostral at a 40° angle to avoid interference with the optical fibers or cranial window. The guide cannula was then anchored to the skull with dental cement (G-CEM ONE neo, GC.dental).

### Striatal local bicuculline injection

Bicuculline methochloride (cat# B7686, Sigma, St. Louis, MO, USA) was dissolved in saline at 100 mg/ml to prepare stock aliquots, which were stored at –30℃. Immediately before each injection, the stock solution was diluted in saline to a final concentration of 0.2 mg/ml.

For single-injection experiments, a small burr hole was made under isoflurane anesthesia, which was discontinued immediately afterward. A disposable glass micropipette filled with bicuculline solution was gently inserted into the dSTR (AP: 0.0 mm, ML: 2.7 mm, DV: –3.5 mm). After recording baseline behavioral data, bicuculline was injected using a glass micropipette attached to a microinjector (I-31, Narishige). The micropipette was left in place until the end of the experiment to minimize tissue damage and prevent leakage into adjacent brain regions.

For repeated-injection experiments (e.g., in DREADD experiments), an injection cannula (AMI-5T or CXMI-8T, Eicom) attached to a microsyringe (cat# N1701, Hamilton, Reno, NV, USA) containing bicuculline solution was inserted into the chronically implanted guide cannula (see above).

In both injection protocols, 0.20 µl of bicuculline was injected over ∼1 min.

### Chemogenetic manipulation

To chemogenetically suppress the activity of specific neurons, the DREADD agonist clozapine-N-oxide (CNO; cat# HY-17366, MedChemExpress, Monmouth Junction, NJ, USA) was administered i.p. to mice expressing hM4Di-mCherry or mCherry (control) in IC neurons or IC-projecting dmTh neurons. CNO was dissolved in dimethyl sulfoxide (DMSO; cat# 13445-74, Nacalai Tesque, Kyoto, Japan) at 100 mg/ml to prepare stock aliquots, which were stored at –30℃. The stock solution was diluted in saline to a final concentration of 2 mg/ml immediately before use.

In the experiments shown in Figures 5, 6, S3, and S4, CNO or vehicle (2% DMSO) was administered at 5 ml/kg i.p (equivalent to 10 mg/kg CNO) 30 min before baseline recordings. Each trial was conducted with an interval of at least 72 h to avoid residual effects of CNO or bicuculline from the previous session.

### EEG and EMG recordings

After the mice were secured in the head-fixation apparatus, EEG and EMG electrodes were connected to an amplifier with an integrated A/D converter (PowerLab 26T, ADInstruments, Dunedin, New Zealand). EEG and EMG signals were sampled at a sampling rate of 10 kHz. Behavioral videos were simultaneously recorded at 100 frames/s with a digital camera (acA800-510um, Basler, Ahrensburg, Germany) positioned in front of the mice.

Baseline recordings were obtained for 5 min before bicuculline injection. Following bicuculline injection, recordings continued for 40 min. The acquired EEG and EMG data were preprocessed in MATLAB (MathWorks, Natick, MA, USA). After downsampling to 1 kHz, the data were bandpass filtered (EEG: 0.5 – 120 Hz; EMG: 5 – 500 Hz) and notch-filtered at 60 Hz to remove line noise.

### Fiber photometry recording

The implanted optical fibers in the ipsilateral M1 or bilateral IC of mice expressing GCaMP6f in these brain regions were connected to a multichannel fiber photometry system (R821, RWD Life Science, Shenzhen, China) via a bundle branching patch cord (BBP(4) 400/430/1100-0.57_2m_FCM-4xZF1.25(F)_LAF, Doric Lenses, Québec, Canada), which enabled simultaneous recording from multiple brain regions. Dual-wavelength recording (470 nm and 410 nm) was performed: the 470-nm LED excitation light was used to detect GCaMP6f fluorescence, whereas the 410-nm one was used to measure calcium-independent isosbestic signals. Neuronal Ca^2+^ signals and behavioral videos were synchronously recorded at 60 Hz. Recordings were conducted for 5 min before and 40 min after bicuculline injection.

Data were analyzed using custom-written MATLAB scripts. Briefly, both 470-nm and 410-nm signals were smoothed using a locally weighted linear regression filter to reduce high-frequency noise. The fitted control signal was obtained by regressing the 410-nm signal against the 470-nm signal using a quantile regression method. ΔF/F was calculated as (F-F_0_)/ F_0_, where F represents the 470-nm signal and F_0_ represents the fitted control signal.

### In vivo two-photon microscopic calcium imaging

Two-photon images were acquired from the mice expressing GCaMP6f in the ipsilateral M1 using an A1MP^+^ system (NIS-Elements, Nikon Instech Co., Ltd, Tokyo, Japan) equipped with a 10× water-immersion objective lens (numerical aperture [NA], 0.3; working distance 3.5 mm; cat# MRH07120, Nikon Instech Co., Ltd) and a mode-locked Ti:sapphire Chameleon Ultra II laser (Chameleon Vision, Coherent, Santa Clara, CA, USA) set at 950 nm.^77^ To quantify neuronal activity, time-lapse images were acquired at a resolution of 512 × 512 pixels (scanning rate, 2 Hz; 2× digital zoom).

Before bicuculline injection, two continuous image series of 1000 frames each were acquired from the ipsilateral M1 at depths of 250 and 500 µm from the cortical surface. After bicuculline injection, continuous 1000-frame images were captured again from the same fields of view. Images were repeatedly acquired for 5-25 min after bicuculline injection. As shown in Figures 1 and S1, this period corresponds to the time window in which tic-like movements were clearly observed.

During two-photon microscopic imaging, behavioral videos were simultaneously recorded at 30 frames/s with an analog infrared-sensitive CCD camera (ITC-405HIR, ITS, Tokyo, Japan).

### Histology

Mice were deeply anesthetized with ketamine/xylazine and transcardially perfused with 4% paraformaldehyde (PFA). The brain was extracted, post-fixed in 4% PFA overnight, and soaked in 15% and then 30% sucrose for 2-3 days for cryoprotection. The brains were sectioned into 50-µm coronal slices on a microtome (OFT5000, Bright Instruments, Cambridgeshire, UK). For c-Fos immunohistochemistry, brain slices were permeabilized with 0.5% Triton X-100 (Cat# 35501-15, Nacalai Tesque) in PBS for 3 h at room temperature. After blocking with 5% bovine serum albumin (BSA; Cat# 01860-36, Nacalai Tesque) for 1 h, the slices were incubated overnight at 4 ℃ with primary antibody against c-Fos (cat# sc-52, Santa Cruz Biotechnology, Dallas, TX, USA; 1:1000) diluted in PBS. After washing in PBS, the slices were incubated overnight at 4℃ with Cy3-conjugated anti-rabbit IgG secondary antibody (cat# 711-165-152, Jackson ImmunoResearch, West Grove, PA; 1:1000) in PBS. The slices were then mounted on glass slides with DAPI-containing mounting medium (H-1200, Vector Laboratories, Newark, CA, USA).

Fluorescence images were captured using a KEYENCE BZ-X810 microscope (KEYENCE, Osaka, Japan) with a 10× objective lens (NA 0.30) or an Olympus FV-3000 confocal microscope (Olympus, Tokyo, Japan) equipped with a 40× objective lens (NA 0.95).

## QUANTIFICATION AND STATISTICAL ANALYSIS

### Behavioral analysis

Tic-like movements were quantified in head-fixed mice following unilateral striatal bicuculline injection. User-defined body parts were tracked using DeepLabCut (https://github.com/DeepLabCut).^78^ A custom-made MATLAB script was used to compute the speed (i.e., scalar velocity) and acceleration of movements in the contralateral wrist or the contralateral corner of the mouth based on the DeepLabCut tracking trajectories. Motor tics were detected based on the timing of spike signals in the ipsilateral M1 cortical activity obtained from EEG or fiber photometry recordings (see below for details), because the M1 spikes were generally accompanied by motor tics (Figures 1, 3, and S1). For EEG-based tic detection, the speed of body movements was calculated within a 0–0.3 s time window after M1 spikes. For fiber photometry-based tic detection, the speed was calculated in a –0.2 to +0.1 s window relative to the M1 spike, as tic-related Ca^2+^ activity typically lagged behind the onset of tic-like movements by ∼0.2 s (see Figure 3).

In Figures 1K and S1, the area under the curve (AUC) of movement speed of the contralateral wrist and mouth was calculated within a 0–0.3 s time window after EEG spikes associated with tics as follows:

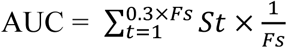

where *St* is the speed of the targeted body part at each time point after an EEG spike, and Fs is the sampling frequency of the behavioral video.

A movement was classified as a motor tic if the speed of the contralateral wrist or corner of the mouth exceeded the mean + 0.5 SD and the maximum acceleration exceeded the mean + 2 SD of the entire recording period. This automated detection method was largely consistent with tic identification by visual inspection.

For DREADD experiments, tic intensity was calculated as (S_wrist_/S1_wrist_ + S_mouth_/S1_mouth_)/2, where S is the average speed of the contralateral wrist or contralateral corner of the mouth during tic expression (0-30 min after bicuculline injection) and S1 is that of the first test (vehicle injection). The relative intensity value of the first test was therefore set to 1.

During *in vivo* two-photon imaging, the resolution of behavioral monitoring with an analog infrared-sensitive CCD camera was insufficient for accurate tracking of mouth-corner movements because of the low light intensity. Therefore, only forelimb trajectories were used to assess tic-like movements. Forelimb tics were automatically detected when their speed exceeded the mean + 0.5 SD and their peak acceleration exceeded the mean + 2 SD. Although this method relied solely on forelimb speed, detected tic onsets were largely consistent with those identified by an experienced observer.

### Spike detection and signal analysis

In the analyses of EEG and fiber photometry data, preprocessed signals (as described in the “EEG and EMG recordings” and “Fiber photometry recording” sections) were converted into robust *Z* scores. Because our data contained numerous spike signals (i.e., outliers), we used a robust z score based on the median and interquartile range (IQR, representing the central 50% of the distribution) to reduce the influence of outliers, in contrast to a conventional mean- and SD-based *Z* score.^79^

The robust *Z* score was calculated as:

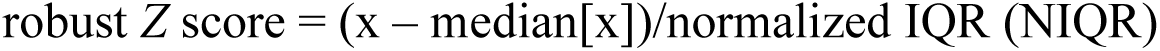

where NIQR was obtained by converting the IQR to the equivalent of 1 SD in a normal distribution by multiplying by a normalization factor of 0.7413. If the data followed a normal distribution, the robust *Z* score coincided with the conventional *Z* score.

Signal peaks with a prominence of at least 1 × NIQR were detected using the “findpeaks” function in MATLAB. The prominence values of the detected peaks were then subjected to k-means clustering to automatically determine the detection threshold. For EEG data, the number of clusters was set to 3; for fiber photometry data, it was set to 4. Peaks classified into the cluster with the lowest prominence were excluded as non-spike events, and the remaining peaks were defined as spike signals.

For calculating the mean amplitude of tic-related Ca^2+^ spikes shown in Figures 3E and 6D, the average robust *Z* score of the prominence values was computed within a 0–0.3 s time window following the onset of tic-like movements.

### EMG analysis

For quantification of EMG amplitude in Figure 1, we calculated an EMG envelope, a smoothed curve that tracks temporal changes in EMG signal amplitude.^80^ To compute the envelope, the preprocessed signals (as described in the “EEG and EMG recordings” section) were rectified and subsequently low-pass filtered at 2 Hz. The envelope values were then normalized using the robust *Z* score, as described in the “Spike detection and signal analysis” section. For the mean EMG amplitude shown in Figure 1J, the average robust Z score of EMG amplitude was calculated within a 0–0.3 s window following EEG spikes associated with tics.

### Image analysis

Images were analyzed using Fiji/ImageJ (National Institutes of Health, Bethesda, MD, USA) and MATLAB.

For counting c-Fos positive cells in Figure 2, images were binarized using the built-in auto-threshold “Triangle” function in ImageJ. After background noise was removed using the “Remove Outliers” function, c-Fos positive cells were automatically counted using the “Analyze Particles” function.

For counting EGFP-labeled thalamic neurons in Figure 4, coronal sections containing target brain regions were selected at 300-µm intervals, and EGFP-positive cells were manually counted. Cell counts from all sections in each thalamic region were summed and normalized to those in the ventromedial (VM) thalamus.

For quantification of tdTomato-labeled axonal projections originating from the substantia nigra pars reticulata (SNr) in Figure S2, images were binarized using the built-in auto-threshold “Triangle” function in ImageJ to separate axonal fluorescence from background. The fraction of suprathreshold pixels was calculated for each thalamic subregion and normalized to the value in VM thalamus.

Two-photon Ca^2+^ images were corrected for focal plane displacement using the “TurboReg” plug-in in ImageJ. To estimate neuronal activity, regions of interest (ROIs) were determined using a semi-automated MATLAB-based toolbox with a graphical user interface (EZcalcium; https://github.com/porteralab/EZcalcium).^81^ A Ca^2+^ event was defined as a fluorescence increase with a peak amplitude exceeding the baseline + 2 SD, where baseline was defined as the 35th percentile of the total fluorescence distribution.^82^ The mean amplitude and frequency of Ca^2+^ events were calculated for each detected neuron.

To assess response probability, we quantified how reliably each neuron responded to tic-like movements. A neuron was considered “responsive” if the fluorescence intensity in the two frames after tic onset exceeded the mean of the two preceding frames by more than 2 SD. Response probability was calculated by dividing the number of responsive events by the total number of tic events.

### Statistics

Statistical analyses were performed using MATLAB and R (version 4.4.2; with the “tidyverse”, “rstatix”, “afex”, “emmeans” packages). Details of statistical tests are provided in the figure legends. Statistical tests included paired *t*-test, one-sample *t*-tests with Holm correction, one-way ANOVA (independent or repeated-measures) followed by pairwise *t*-tests with Holm correction, and Kruskal-Wallis test followed by Dunn’s multiple comparison tests with Holm correction.

We did not routinely perform formal tests for normality, as their results are highly sensitive to sample size. When the sample size was large (n > 30), *t*-tests are generally robust to moderate deviations from normality. In such cases, we visually inspected the data distribution and applied parametric tests unless the data exhibited extreme skewness or kurtosis. When the sample size was small, we considered reliable estimation of the population distribution difficult, and we generally used parametric tests unless there was a clear reason to assume non-normality.

Data analyzed with parametric tests are presented as mean ± SEM, while those analyzed with non-parametric tests are shown as box plots. Box plots indicate the 25th and 75th percentiles, the median, and whiskers extending to data within a 1.5 x IQR. All tests were two-tailed with significance set at α = 0.05.

## Notes

### Competing Interest Statement

The authors have declared no competing interest.

### Summary of Updates

Summary revised; Supplementary Figure S2 removed; Supplementary Figures S4-S5 changed; Funding Information updated; other minor corrections

## REFERENCES

1. Diagnostic and statistical manual of mental disorders : DSM-5-TR. (2022). (American Psychiatric Association Publishing). 10.1176/appi.books.9780890425787.

2. Bullen, J.G., and Hemsley, D.R. (1983). Sensory experience as a trigger in Gilles de la Tourette’s syndrome. J Behav Ther Exp Psychiatry 14, 197–201. 10.1016/0005-7916(83)90048-4.

3. Kurlan, R., Lichter, D., and Hewitt, D. (1989). Sensory tics in Tourette’s syndrome. Neurology 39, 731–734. 10.1212/wnl.39.5.731.

4. Leckman, J.F., Walker, D.E., and Cohen, D.J. (1993). Premonitory urges in Tourette’s syndrome. Am J Psychiatry 150, 98–102. 10.1176/ajp.150.1.98.

5. Houghton, D.C., Capriotti, M.R., Conelea, C.A., and Woods, D.W. (2014). Sensory Phenomena in Tourette Syndrome: Their Role in Symptom Formation and Treatment. Curr Dev Disord Rep 1, 245–251. 10.1007/s40474-014-0026-2.

6. Jankovic, J. (2001). Tourette’s syndrome. N Engl J Med 345, 1184–1192. 10.1056/NEJMra010032.

7. Leckman, J.F. (2002). Tourette’s syndrome. Lancet 360, 1577–1586. 10.1016/S0140-6736(02)11526-1.

8. Johnson, K.A., Worbe, Y., Foote, K.D., Butson, C.R., Gunduz, A., and Okun, M.S. (2023). Tourette syndrome: clinical features, pathophysiology, and treatment. Lancet Neurol 22, 147–158. 10.1016/S1474-4422(22)00303-9.

9. Robertson, M.M. (2008). The prevalence and epidemiology of Gilles de la Tourette syndrome. Part 1: the epidemiological and prevalence studies. J Psychosom Res 65, 461–472. 10.1016/j.jpsychores.2008.03.006.

10. Knight, T., Steeves, T., Day, L., Lowerison, M., Jette, N., and Pringsheim, T. (2012). Prevalence of tic disorders: a systematic review and meta-analysis. Pediatr Neurol 47, 77–90. 10.1016/j.pediatrneurol.2012.05.002.

11. Scharf, J.M., Miller, L.L., Gauvin, C.A., Alabiso, J., Mathews, C.A., and Ben-Shlomo, Y. (2015). Population prevalence of Tourette syndrome: a systematic review and meta-analysis. Mov Disord 30, 221–228. 10.1002/mds.26089.

12. Jafari, F., Abbasi, P., Rahmati, M., Hodhodi, T., and Kazeminia, M. (2022). Systematic Review and Meta-Analysis of Tourette Syndrome Prevalence; 1986 to 2022. Pediatr Neurol 137, 6–16. 10.1016/j.pediatrneurol.2022.08.010.

13. Seideman, M.F., and Seideman, T.A. (2020). A Review of the Current Treatment of Tourette Syndrome. J Pediatr Pharmacol Ther 25, 401–412. 10.5863/1551-6776-25.5.401.

14. Yael, D., Israelashvili, M., and Bar-Gad, I. (2016). Animal Models of Tourette Syndrome-From Proliferation to Standardization. Front Neurosci 10, 132. 10.3389/fnins.2016.00132.

15. Tarsy, D., Pycock, C.J., Meldrum, B.S., and Marsden, C.D. (1978). Focal contralateral myoclonus produced by inhibition of GABA action in the caudate nucleus of rats. Brain 101, 143–162. 10.1093/brain/101.1.143.

16. Bronfeld, M., Yael, D., Belelovsky, K., and Bar-Gad, I. (2013). Motor tics evoked by striatal disinhibition in the rat. Front Syst Neurosci 7, 50. 10.3389/fnsys.2013.00050.

17. Crossman, A.R., Mitchell, I.J., Sambrook, M.A., and Jackson, A. (1988). Chorea and myoclonus in the monkey induced by gamma-aminobutyric acid antagonism in the lentiform complex. The site of drug action and a hypothesis for the neural mechanisms of chorea. Brain 111 ( *Pt 5*), 1211–1233. 10.1093/brain/111.5.1211.

18. McCairn, K.W., Bronfeld, M., Belelovsky, K., and Bar-Gad, I. (2009). The neurophysiological correlates of motor tics following focal striatal disinhibition. Brain 132, 2125–2138. 10.1093/brain/awp142.

19. Worbe, Y., Baup, N., Grabli, D., Chaigneau, M., Mounayar, S., McCairn, K., Féger, J., and Tremblay, L. (2009). Behavioral and movement disorders induced by local inhibitory dysfunction in primate striatum. Cereb Cortex 19, 1844–1856. 10.1093/cercor/bhn214.

20. Kawaguchi, Y., Wilson, C.J., Augood, S.J., and Emson, P.C. (1995). Striatal interneurones: chemical, physiological and morphological characterization. Trends Neurosci 18, 527–535. 10.1016/0166-2236(95)98374-8.

21. Kalanithi, P.S., Zheng, W., Kataoka, Y., DiFiglia, M., Grantz, H., Saper, C.B., Schwartz, M.L., Leckman, J.F., and Vaccarino, F.M. (2005). Altered parvalbumin-positive neuron distribution in basal ganglia of individuals with Tourette syndrome. Proc Natl Acad Sci U S A 102, 13307–13312. 10.1073/pnas.0502624102.

22. Kataoka, Y., Kalanithi, P.S., Grantz, H., Schwartz, M.L., Saper, C., Leckman, J.F., and Vaccarino, F.M. (2010). Decreased number of parvalbumin and cholinergic interneurons in the striatum of individuals with Tourette syndrome. J Comp Neurol 518, 277–291. 10.1002/cne.22206.

23. Shapiro, E., Shapiro, A.K., Fulop, G., Hubbard, M., Mandeli, J., Nordlie, J., and Phillips, R.A. (1989). Controlled study of haloperidol, pimozide and placebo for the treatment of Gilles de la Tourette’s syndrome. Arch Gen Psychiatry 46, 722–730. 10.1001/archpsyc.1989.01810080052006.

24. Kelley, A.E., Lang, C.G., and Gauthier, A.M. (1988). Induction of oral stereotypy following amphetamine microinjection into a discrete subregion of the striatum. Psychopharmacology (Berl) 95, 556–559. 10.1007/BF00172976.

25. Conceição, V.A., Dias, Â., Farinha, A.C., and Maia, T.V. (2017). Premonitory urges and tics in Tourette syndrome: computational mechanisms and neural correlates. Curr Opin Neurobiol 46, 187–199. 10.1016/j.conb.2017.08.009.

26. Jackson, S.R., Loayza, J., Crighton, M., Sigurdsson, H.P., Dyke, K., and Jackson, G.M. (2020). The role of the insula in the generation of motor tics and the experience of the premonitory urge-to-tic in Tourette syndrome. Cortex 126, 119–133. 10.1016/j.cortex.2019.12.021.

27. Bohlhalter, S., Goldfine, A., Matteson, S., Garraux, G., Hanakawa, T., Kansaku, K., Wurzman, R., and Hallett, M. (2006). Neural correlates of tic generation in Tourette syndrome: an event-related functional MRI study. Brain 129, 2029–2037. 10.1093/brain/awl050.

28. Lerner, A., Bagic, A., Boudreau, E.A., Hanakawa, T., Pagan, F., Mari, Z., Bara-Jimenez, W., Aksu, M., Garraux, G., Simmons, J.M., et al. (2007). Neuroimaging of neuronal circuits involved in tic generation in patients with Tourette syndrome. Neurology 68, 1979–1987. 10.1212/01.wnl.0000264417.18604.12.

29. Lee, J., Wang, W., and Sabatini, B.L. (2020). Anatomically segregated basal ganglia pathways allow parallel behavioral modulation. Nat Neurosci 23, 1388–1398. 10.1038/s41593-020-00712-5.

30. Foster, N.N., Barry, J., Korobkova, L., Garcia, L., Gao, L., Becerra, M., Sherafat, Y., Peng, B., Li, X., Choi, J.H., et al. (2021). The mouse cortico-basal ganglia-thalamic network. Nature 598, 188–194. 10.1038/s41586-021-03993-3.

31. McElvain, L.E., Chen, Y., Moore, J.D., Brigidi, G.S., Bloodgood, B.L., Lim, B.K., Costa, R.M., and Kleinfeld, D. (2021). Specific populations of basal ganglia output neurons target distinct brain stem areas while collateralizing throughout the diencephalon. Neuron 109, 1721–1738.e1724. 10.1016/j.neuron.2021.03.017.

32. Aoki, S., Smith, J.B., Li, H., Yan, X., Igarashi, M., Coulon, P., Wickens, J.R., Ruigrok, T.J., and Jin, X. (2019). An open cortico-basal ganglia loop allows limbic control over motor output via the nigrothalamic pathway. Elife 8. 10.7554/eLife.49995.

33. Hariz, M.I., and Robertson, M.M. (2010). Gilles de la Tourette syndrome and deep brain stimulation. Eur J Neurosci 32, 1128–1134. 10.1111/j.1460-9568.2010.07415.x.

34. 34. Arnts, H., Coolen, S.E., Fernandes, F.W., Schuurman, R., Krauss, J.K., Groenewegen, H.J., and van den Munckhof, P. (2023). The intralaminar thalamus: a review of its role as a target in functional neurosurgery. Brain Commun 5, fcad003. 10.1093/braincomms/fcad003.

35. Manvich, D.F., Webster, K.A., Foster, S.L., Farrell, M.S., Ritchie, J.C., Porter, J.H., and Weinshenker, D. (2018). The DREADD agonist clozapine N-oxide (CNO) is reverse-metabolized to clozapine and produces clozapine-like interoceptive stimulus effects in rats and mice. Sci Rep 8, 3840. 10.1038/s41598-018-22116-z.

36. Rodd, Z.A., Engleman, E.A., Truitt, W.A., Burke, A.R., Molosh, A.I., Bell, R.L., and Hauser, S.R. (2022). CNO Administration Increases Dopamine and Glutamate in the Medial Prefrontal Cortex of Wistar Rats: Further Concerns for the Validity of the CNO-activated DREADD Procedure. Neuroscience 491, 176–184. 10.1016/j.neuroscience.2022.03.028.

37. Scharf, J.M., Yu, D., Mathews, C.A., Neale, B.M., Stewart, S.E., Fagerness, J.A., Evans, P., Gamazon, E., Edlund, C.K., Service, S.K., et al. (2013). Genome-wide association study of Tourette’s syndrome. Mol Psychiatry 18, 721–728. 10.1038/mp.2012.69.

38. Willsey, A.J., Fernandez, T.V., Yu, D., King, R.A., Dietrich, A., Xing, J., Sanders, S.J., Mandell, J.D., Huang, A.Y., Richer, P., et al. (2017). De Novo Coding Variants Are Strongly Associated with Tourette Disorder. Neuron 94, 486–499.e489. 10.1016/j.neuron.2017.04.024.

39. Lu, Q., Zhou, Y., Qian, Q., Chen, Z., Tan, Q., Chen, H., Yin, F., Wang, Y., Liu, Z., Tian, P., and Sun, D. (2024). Whole-exome sequencing identifies high-confidence genes for tic disorders in a Chinese Han population. Clin Chim Acta 561, 119759. 10.1016/j.cca.2024.119759.

40. Kowalski, T.F., Wang, R., and Tischfield, M.A. (2025). Genetic advances and translational phenotypes in rodent models for Tourette disorder. Curr Opin Neurobiol 90, 102967. 10.1016/j.conb.2024.102967.

41. Godar, S.C., Mosher, L.J., Di Giovanni, G., and Bortolato, M. (2014). Animal models of tic disorders: a translational perspective. J Neurosci Methods 238, 54–69. 10.1016/j.jneumeth.2014.09.008.

42. McCairn, K.W., Nagai, Y., Hori, Y., Ninomiya, T., Kikuchi, E., Lee, J.Y., Suhara, T., Iriki, A., Minamimoto, T., Takada, M., et al. (2016). A Primary Role for Nucleus Accumbens and Related Limbic Network in Vocal Tics. Neuron 89, 300–307. 10.1016/j.neuron.2015.12.025.

43. Gittis, A.H., Berke, J.D., Bevan, M.D., Chan, C.S., Mallet, N., Morrow, M.M., and Schmidt, R. (2014). New roles for the external globus pallidus in basal ganglia circuits and behavior. J Neurosci 34, 15178–15183. 10.1523/JNEUROSCI.3252-14.2014.

44. Matsumura, M., Tremblay, L., Richard, H., and Filion, M. (1995). Activity of pallidal neurons in the monkey during dyskinesia induced by injection of bicuculline in the external pallidum. Neuroscience 65, 59–70. 10.1016/0306-4522(94)00484-m.

45. Grabli, D., McCairn, K., Hirsch, E.C., Agid, Y., Féger, J., François, C., and Tremblay, L. (2004). Behavioural disorders induced by external globus pallidus dysfunction in primates: I. Behavioural study. Brain 127, 2039–2054. 10.1093/brain/awh220.

46. Bronfeld, M., Belelovsky, K., Erez, Y., Bugaysen, J., Korngreen, A., and Bar-Gad, I. (2010). Bicuculline-induced chorea manifests in focal rather than globalized abnormalities in the activation of the external and internal globus pallidus. J Neurophysiol 104, 3261–3275. 10.1152/jn.00093.2010.

47. Tachibana, Y., Kita, H., Chiken, S., Takada, M., and Nambu, A. (2008). Motor cortical control of internal pallidal activity through glutamatergic and GABAergic inputs in awake monkeys. Eur J Neurosci 27, 238–253. 10.1111/j.1460-9568.2007.05990.x.

48. McCairn, K.W., Iriki, A., and Isoda, M. (2013). Global dysrhythmia of cerebro-basal ganglia-cerebellar networks underlies motor tics following striatal disinhibition. J Neurosci 33, 697–708. 10.1523/JNEUROSCI.4018-12.2013.

49. Murakami, J., Tachibana, Y., Akiyama, S., Kato, T., Taniguchi, A., Nakajima, Y., Shimoda, M., Wake, H., Kano, Y., Takada, M., et al. (2019). Oral splint ameliorates tic symptoms in patients with tourette syndrome. Mov Disord 34, 1577–1578. 10.1002/mds.27819.

50. Yoshida, A., Fujio, T., Sato, F., Ali, M.S.S., Haque, T., Ohara, H., Moritani, M., Kato, T., Dostrovsky, J.O., and Tachibana, Y. (2017). Orofacial proprioceptive thalamus of the rat. Brain Struct Funct 222, 2655–2669. 10.1007/s00429-016-1363-1.

51. Sato, F., Uemura, Y., Kanno, C., Tsutsumi, Y., Tomita, A., Oka, A., Kato, T., Uchino, K., Murakami, J., Haque, T., et al. (2017). Thalamo-insular pathway conveying orofacial muscle proprioception in the rat. Neuroscience 365, 158–178. 10.1016/j.neuroscience.2017.09.050.

52. Craig, A.D. (2002). How do you feel? Interoception: the sense of the physiological condition of the body. Nat Rev Neurosci 3, 655–666. 10.1038/nrn894.

53. Craig, A.D. (2009). How do you feel--now? The anterior insula and human awareness. Nat Rev Neurosci 10, 59–70. 10.1038/nrn2555.

54. Namkung, H., Kim, S.H., and Sawa, A. (2017). The Insula: An Underestimated Brain Area in Clinical Neuroscience, Psychiatry, and Neurology. Trends Neurosci 40, 200–207. 10.1016/j.tins.2017.02.002.

55. Bergman, H., Wichmann, T., and DeLong, M.R. (1990). Reversal of experimental parkinsonism by lesions of the subthalamic nucleus. Science 249, 1436–1438. 10.1126/science.2402638.

56. Lozano, A.M., Lang, A.E., Galvez-Jimenez, N., Miyasaki, J., Duff, J., Hutchinson, W.D., and Dostrovsky, J.O. (1995). Effect of GPi pallidotomy on motor function in Parkinson’s disease. Lancet 346, 1383–1387. 10.1016/s0140-6736(95)92404-3.

57. Lang, A.E., and Lozano, A.M. (1998). Parkinson’s disease. First of two parts. N Engl J Med 339, 1044–1053. 10.1056/NEJM199810083391506.

58. Boraud, T., Bezard, E., Bioulac, B., and Gross, C. (1996). High frequency stimulation of the internal Globus Pallidus (GPi) simultaneously improves parkinsonian symptoms and reduces the firing frequency of GPi neurons in the MPTP-treated monkey. Neurosci Lett 215, 17–20. 10.1016/s0304-3940(96)12943-8.

59. Obeso, J.A., Olanow, C.W., Rodriguez-Oroz, M.C., Krack, P., Kumar, R., Lang, A.E., and Group, D.-B.S.f.P.s.D.S. (2001). Deep-brain stimulation of the subthalamic nucleus or the pars interna of the globus pallidus in Parkinson’s disease. N Engl J Med 345, 956–963. 10.1056/NEJMoa000827.

60. Lozano, A.M., Lipsman, N., Bergman, H., Brown, P., Chabardes, S., Chang, J.W., Matthews, K., McIntyre, C.C., Schlaepfer, T.E., Schulder, M., et al. (2019). Deep brain stimulation: current challenges and future directions. Nat Rev Neurol 15, 148–160. 10.1038/s41582-018-0128-2.

61. Vandewalle, V., van der Linden, C., Groenewegen, H.J., and Caemaert, J. (1999). Stereotactic treatment of Gilles de la Tourette syndrome by high frequency stimulation of thalamus. Lancet 353, 724. 10.1016/s0140-6736(98)05964-9.

62. Singer, H.S. (2005). Tourette’s syndrome: from behaviour to biology. Lancet Neurol 4, 149–159. 10.1016/S1474-4422(05)01012-4.

63. Sadikot, A.F., and Rymar, V.V. (2009). The primate centromedian-parafascicular complex: anatomical organization with a note on neuromodulation. Brain Res Bull 78, 122–130. 10.1016/j.brainresbull.2008.09.016.

64. Cover, K.K., and Mathur, B.N. (2021). Rostral Intralaminar Thalamus Engagement in Cognition and Behavior. Front Behav Neurosci 15, 652764. 10.3389/fnbeh.2021.652764.

65. Alexander, G.E., DeLong, M.R., and Strick, P.L. (1986). Parallel organization of functionally segregated circuits linking basal ganglia and cortex. Annu Rev Neurosci 9, 357–381. 10.1146/annurev.ne.09.030186.002041.

66. Alexander, G.E., and Crutcher, M.D. (1990). Functional architecture of basal ganglia circuits: neural substrates of parallel processing. Trends Neurosci 13, 266–271. 10.1016/0166-2236(90)90107-l.

67. Tremblay, L., Worbe, Y., Thobois, S., Sgambato-Faure, V., and Féger, J. (2015). Selective dysfunction of basal ganglia subterritories: From movement to behavioral disorders. Mov Disord 30, 1155–1170. 10.1002/mds.26199.

68. Papez, J. (1937). A proposed mechanism of emotion. Archives of Neurology and Psychiatry 38, 725–743. 10.1001/archneurpsyc.1937.02260220069003.

69. Saper, C.B. (1982). Convergence of autonomic and limbic connections in the insular cortex of the rat. J Comp Neurol 210, 163–173. 10.1002/cne.902100207.

70. Fudge, J.L., Breitbart, M.A., Danish, M., and Pannoni, V. (2005). Insular and gustatory inputs to the caudal ventral striatum in primates. J Comp Neurol 490, 101–118. 10.1002/cne.20660.

71. Gehrlach, D.A., Weiand, C., Gaitanos, T.N., Cho, E., Klein, A.S., Hennrich, A.A., Conzelmann, K.K., and Gogolla, N. (2020). A whole-brain connectivity map of mouse insular cortex. Elife 9. 10.7554/eLife.55585.

72. Tyler, W.J., Lani, S.W., and Hwang, G.M. (2018). Ultrasonic modulation of neural circuit activity. Curr Opin Neurobiol 50, 222–231. 10.1016/j.conb.2018.04.011.

73. Rabut, C., Yoo, S., Hurt, R.C., Jin, Z., Li, H., Guo, H., Ling, B., and Shapiro, M.G. (2020). Ultrasound Technologies for Imaging and Modulating Neural Activity. Neuron 108, 93–110. 10.1016/j.neuron.2020.09.003.

74. Martínez-Fernández, R., Paschen, S., Del Álamo, M., Rodríguez-Rojas, R., Pineda-Pardo, J.A., Blesa, J., Kaplitt, M.G., Deuschl, G., and Obeso, J.A. (2025). Focused ultrasound therapy for movement disorders. Lancet Neurol 24, 698–712. 10.1016/S1474-4422(25)00210-8.

75. Tanahira, C., Higo, S., Watanabe, K., Tomioka, R., Ebihara, S., Kaneko, T., and Tamamaki, N. (2009). Parvalbumin neurons in the forebrain as revealed by parvalbumin-Cre transgenic mice. Neurosci Res 63, 213–223. 10.1016/j.neures.2008.12.007.

76. Franklin, K.B.J., and Paxinos, G. (2008). The Mouse Brain in Stereotaxic Coordinates, Compact, 3rd Edition.

77. Sotani, N., Kusuhara, S., Nishisho, R., Kuno, H., Shima, H., Haruwaka, K., Mori, Y., Kishi, M., Furuyashiki, T., Kobayashi, K., et al. Transpupillary in vivo two-photon imaging reveals enhanced surveillance of retinal microglia in diabetic mice (in press). Proc Natl Acad Sci.

78. Mathis, A., Mamidanna, P., Cury, K.M., Abe, T., Murthy, V.N., Mathis, M.W., and Bethge, M. (2018). DeepLabCut: markerless pose estimation of user-defined body parts with deep learning. Nat Neurosci 21, 1281–1289. 10.1038/s41593-018-0209-y.

79. Kawanabe-Kobayashi, R., Uchiyama, S., Yoshihara, K., Kojima, D., McHugh, T., Hatada, I., Matsui, K., Tanaka, K.F., and Tsuda, M. (2025). Descending locus coeruleus noradrenergic signaling to spinal astrocyte subset is required for stress-induced pain facilitation (Preprint). Elife. 10.7554/elife.104453.1.

80. Spulák, D., Cmejla, R., Bačáková, R., Kračmar, B., Satrapová, L., and Novotný, P. (2014). Muscle activity detection in electromyograms recorded during periodic movements. Comput Biol Med 47, 93–103. 10.1016/j.compbiomed.2014.01.013.

81. Cantu, D.A., Wang, B., Gongwer, M.W., He, C.X., Goel, A., Suresh, A., Kourdougli, N., Arroyo, E.D., Zeiger, W., and Portera-Cailliau, C. (2020). EZcalcium: Open-Source Toolbox for Analysis of Calcium Imaging Data. Front Neural Circuits 14, 25. 10.3389/fncir.2020.00025.

82. Okada, T., Kato, D., Nomura, Y., Obata, N., Quan, X., Morinaga, A., Yano, H., Guo, Z., Aoyama, Y., Tachibana, Y., et al. (2021). Pain induces stable, active microcircuits in the somatosensory cortex that provide a therapeutic target. Sci Adv 7. 10.1126/sciadv.abd8261.

