## Supplementary material for "Thalamo-insular pathway regulates tic generation via motor-limbic crosstalk"

**Key Resources Table**

**Supplemental Figures**

Figure S1: Orofacial tic-like movements induced by striatal disinhibition, related to Figure 1.

Figure S2: Quantification of SNr-driven axon terminals across thalamic subregions, related to Figure 4.

Figure S3: Chemogenetic inhibition of the thalamo-insular pathway reduces tic intensity without affecting tic frequency, related to Figure 6.

Figure S4: Chemogenetic inhibition of IC or the dmTh-IC pathway alters M1 activity in a layer-specific manner, related to Figure 6.

Figure S5. Motor-limbic crosstalk underlying tic generation, related to Figure 6.

**Supplemental Videos**

Video S1: Tic-like movements and tic-related EEG spikes induced by striatal bicuculline injection, related to Figure 1.

Video S2: Suppression of motor tics by chemogenetic inhibition of IC or the dmTh-IC pathway, related to Figure 5.

Video S3: Synchronized neuronal activity in M1 following striatal bicuculline injection, related to Figure 6.

### Key Resources Table

| REAGENT or RESOURCE | SOURCE | IDENTIFIER |
|---|---|---|
| Antibodies | | |
| Rabbit anti-c-Fos | Santa Cruz Biotechnology | Cat# sc-52 |
| Donkey anti-rabbit IgG H&L (Cy3) | Jackson ImmunoResearch | Cat# 711-165-152 |
| Bacterial and viral strains | | |
| AAV8-hSyn-hM4Di-mCherry | Addgene | Cat# 50475-AAV8 |
| AAV8-hSyn-mCherry | Kobayashi Lab | N/A |
| AAVDJ-EF1α-DIO-hM4Di-mCherry | Kobayashi Lab | N/A |
| AAVDJ-EF1α-DIO-mCherry | Kobayashi Lab | N/A |
| AAV2retro-CAGGS-Cre | Kobayashi Lab | N/A |
| AAV1-Syn-GCaMP6f-WPRE-SV40 | Addgene | Cat# 100835-AAV1 |
| AAV1-CAG-tdTomato | Addgene | Cat# 59462-AAV1 |
| AAV1-CAG-Flex-tdTomato | Addgene | Cat# 28306-AAV1 |
| AAV2retro-CAGGS-EGFP | Kobayashi Lab | N/A |
| Chemicals, peptides, and recombinant proteins | | |
| UNIFAST Ⅱ | GC.dental | N/A |
| G-CEM ONE neo | GC.dental | N/A |
| G-CEM ONE adhesive-enhancing primer | GC.dental | N/A |
| Fuji Lute | GC.dental | N/A |
| Super Bond | Sun Medical | N/A |
| Vetbond | 3M Co. | N/A |
| NOR-61 | Norland Products | N/A |
| Clozapine N-oxide (CNO) | MedChemExpress | Cat# HY-17366, CAS: 34233-69-7 |
| dimethyl sulfoxide (DMSO) | Nacalai tesque | Cat# 13445-74, CAS: 67-68-5 |
| Bicuculline methochloride | Sigma | Cat# B7686, CAS: 38641-83-7 |
| Bovaine Serum Albumin | Nacalai tesque | Cat# 01860-36, CAS: 9048-46-8 |
| Triton X-100 | Nacalai tesque | Cat# 35501-15, CAS: 9002-93-1 |
| VECTASHIELD Mounting Medium for Fluorescence with DAPI | Vector Laboratories | Cat# H-1200 |
| Ketamine | Daiichi Sankyo | CAS: 1867-66-9 |
| Xylazine | Elanco Japan | CAS: 23076-35-9 |
| Isoflurane | Viatris | CAS: 26675-46-7 |

| | | |
|---|---|---|
| Xylocaine DENTAL | Dentsply Sirona | N/A |
| Deposited data | | |
| Source data | This paper | Mendeley Data: https://data.mendeley.com/preview/spv83brm5d?a=884e006d-2eec-41d1-893b-e30892663a09 |
| Experimental models: Organisms/strains | | |
| Mouse: C57BL6/J | Japan SLC | C57BL/6JJmsSlc RRID: MGI: 5488963 |
| Mouse: Tg(Pvalb-cre)1Tama | C. Tanahira et al.[1] | RRID: MGI: 5311448 |
| Software and algorithms | | |
| Matlab | MathWorks | RRID: SCR_001622 |
| EZcalcium | DA Cantu et al.[2] | https://github.com/porteralab/EZcalcium; RRID:SCR_022354 |
| ImageJ | National Institute of Health | RRID: SCR_003070 |
| Fiji (Fiji is just ImageJ) | Opensource | RRID:SCR_002285 |
| DEEPLABCUT | Mathis et al.[3] | https://github.com/DeepLabCut; RRID:SCR_021391 |
| R v4.4.2 | R Core Team | RRID: SCR_001905 |
| CorelDRAW Graphics Suite 2024 | Corel Corporation | RRID: SCR_014235 |
| Other | | |
| 5 mm guide cannula | Eicom | Cat# AG-5T |
| 8 mm guide cannula | Eicom | Cat# CXG-8T |
| 5 mm injection cannula | Eicom | Cat# AMI-5T |
| 8 mm injection cannula | Eicom | Cat# CXMI-8T |
| Microliter syringe | Hamilton | Cat# N1701 |
| PFA-coated stainless-steel wires | A-M SYSTEMS | Cat# 793200 |
| Bundle branching patch cord | Doric Lenses | Ordering code: BBP(4)_400/430/1100-0.57_2m_FCM-4xZF1.25(F)_LAF |
| Optical fiber | RWD Life Science | Cat# R-FOC-BL400C-50NA |

### References Cited:


1. Tanahira, C., Higo, S., Watanabe, K., Tomioka, R., Ebihara, S., Kaneko, T., and Tamamaki, N. (2009). Parvalbumin neurons in the forebrain as revealed by parvalbumin-Cre transgenic mice.

Neurosci Res *63*, 213-223. 10.1016/j.neures.2008.12.007.

2. Cantu, D.A., Wang, B., Gongwer, M.W., He, C.X., Goel, A., Suresh, A., Kourdougli, N., Arroyo, E.D., Zeiger, W., and Portera-Cailliau, C. (2020). EZcalcium: Open-Source Toolbox for Analysis of Calcium Imaging Data. Front Neural Circuits *14*, 25. 10.3389/fncir.2020.00025.
3. Mathis, A., Mamidanna, P., Cury, K.M., Abe, T., Murthy, V.N., Mathis, M.W., and Bethge, M. (2018). DeepLabCut: markerless pose estimation of user-defined body parts with deep learning. Nat Neurosci *21*, 1281-1289. 10.1038/s41593-018-0209-y.

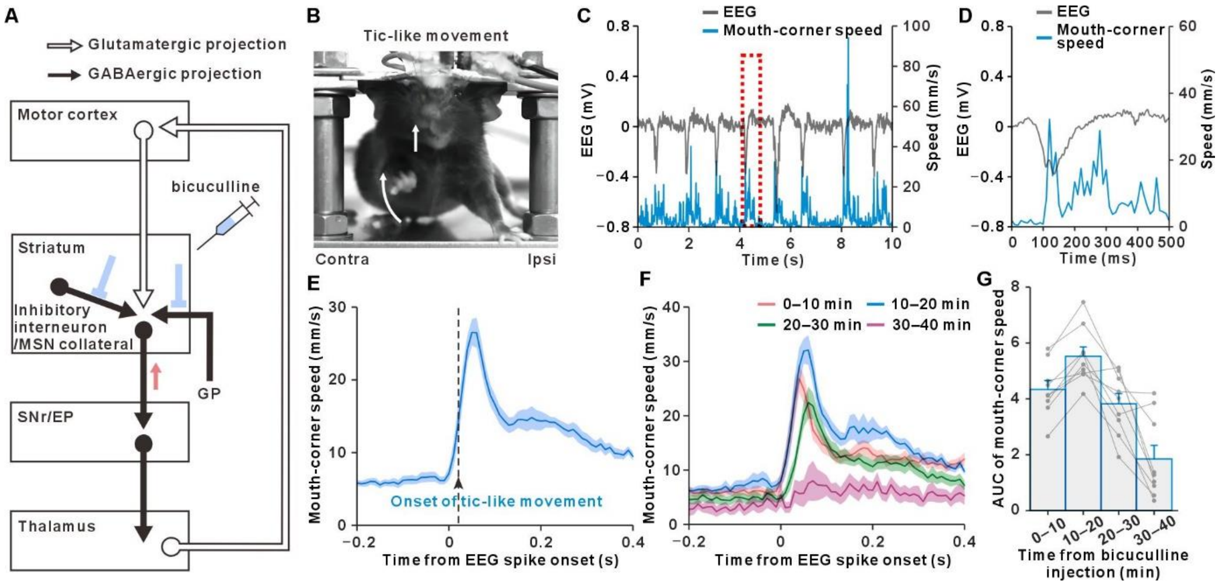


**Figure S1. Orofacial tic-like movements induced by striatal disinhibition. Related to Figure 1.**

(A) Schematic of the cortico-basal ganglia-thalamo-cortical (CBGTC) circuitry illustrating glutamatergic (white arrows) and GABAergic (black arrows) projections, with the site of unilateral bicuculline injection into the striatum indicated. Bicuculline injection inhibits GABAergic inputs to striatal projection neurons. EP, entopeduncular nucleus; GP, globus pallidus; MSN, medium spiny neuron; SNr, substantia nigra pars reticulata.

(B) Image of a head-fixed mouse exhibiting tic-like movements of the mouth (straight arrow) and forelimb (curved arrow) contralateral to the site of bicuculline injection.

(C and D) Representative traces of electroencephalogram (EEG) from the ipsilateral primary motor cortex (M1; black) and mouth-corner movement speed (blue) following bicuculline injection. The region outlined by the red dashed rectangle in (C) is shown at higher magnification in (D).

(E) Averaged traces of mouth-corner movement speed (blue) aligned to the onsets of EEG spikes recorded in the ipsilateral M1 during the 40-min period following bicuculline injection (n = 9 mice). The black arrow indicates the mean latency of tic onsets relative to EEG spike onsets. The shaded area represents ± SEM.

(F) Averaged traces of mouth-corner movement speed aligned to the onsets of EEG spikes, plotted separately for successive 10-min epochs following bicuculline injection (n = 9 mice).

(G) Time course of the area under the curve (AUC) of mouth-corner movement speed within a 0–0.3 s time window following EEG spikes associated with tics, plotted in 10-min bins after bicuculline injection (n = 9 mice). Data are presented as mean ± SEM; individual mouse data are shown in gray.

See also Video S1.

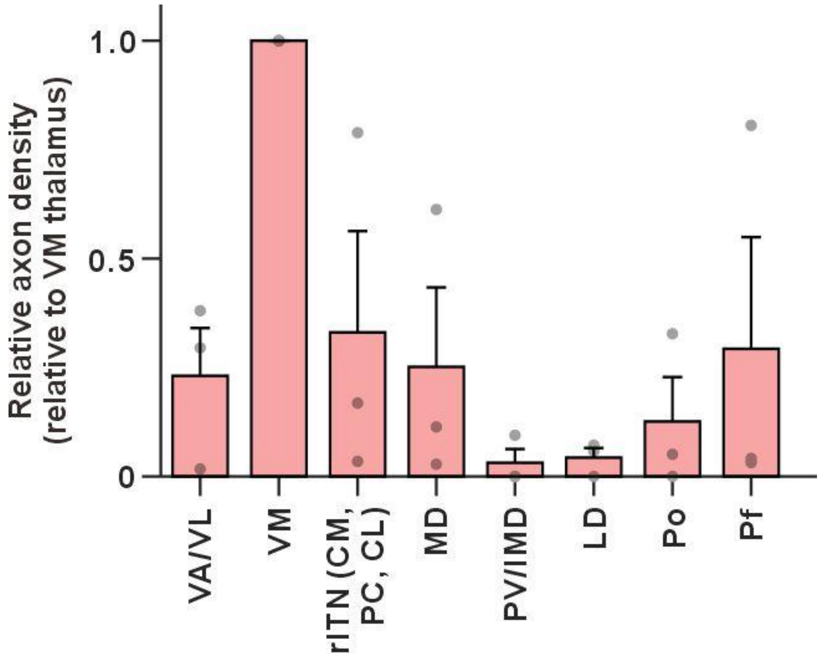


**Figure S2. Quantification of SNr-driven axon terminals across thalamic subregions. Related to Figure 4.**

The fraction of suprathreshold fluorescent pixels from tdTomato-labeled axons originating from parvalbumin (PV)-positive neurons in the substantia nigra pars reticulata (SNr) of PV-Cre mice (shown in Figure 4D) was quantified across multiple thalamic subregions: ventral anterior/ventrolateral (VA/VL) thalamic nuclei, ventromedial (VM) nucleus, rostral intralaminar thalamic nuclei (rITN; central medial [CM], paracentral [PC], and centrolateral [CL] thalamic nuclei), mediodorsal (MD) thalamic nucleus, paraventricular/intermediodorsal (PV/IMD) thalamic nuclei, laterodorsal (LD) thalamic nucleus, posterior (Po) thalamic nucleus, parafascicular (Pf) thalamic nucleus. Data are normalized to values in VM thalamus. Bars indicate mean ± SEM (n = 3 mice); individual data points are shown in gray.

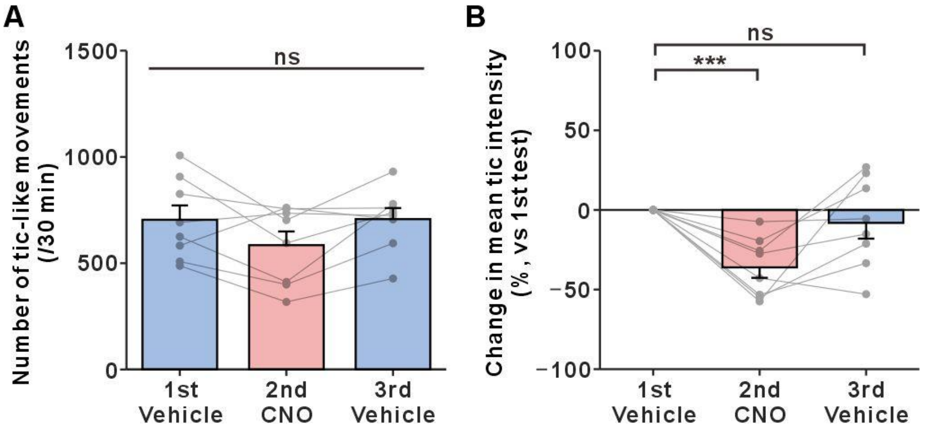


**Figure S3. Chemogenetic inhibition of the thalamo-insular pathway reduces tic intensity without affecting tic frequency. Related to Figure 6.**

(A and B) Effects of chemogenetic inhibition of the dorsomedial thalamus (dmTh) neurons projecting to the insular cortex (IC). Mice received repeated unilateral bicuculline injections into the dSTR via a chronically implanted guide cannula following intraperitoneal administration of vehicle (1st test), clozapine-N-oxide (CNO, 10 mg/kg; 2nd test), and vehicle (3rd test). Tic frequency (A) was unchanged, while tic intensity (B) was significantly reduced during the 30-min period following bicuculline injection (n = 8 mice). Data are presented as mean ± SEM; individual mouse data are shown in gray. Statistical analyses were performed using one-way repeated measures ANOVA (A) or one-sample *t*-tests with Holm correction (B). ns, not significant; ***$p < 0.001$.

See also Video S2.

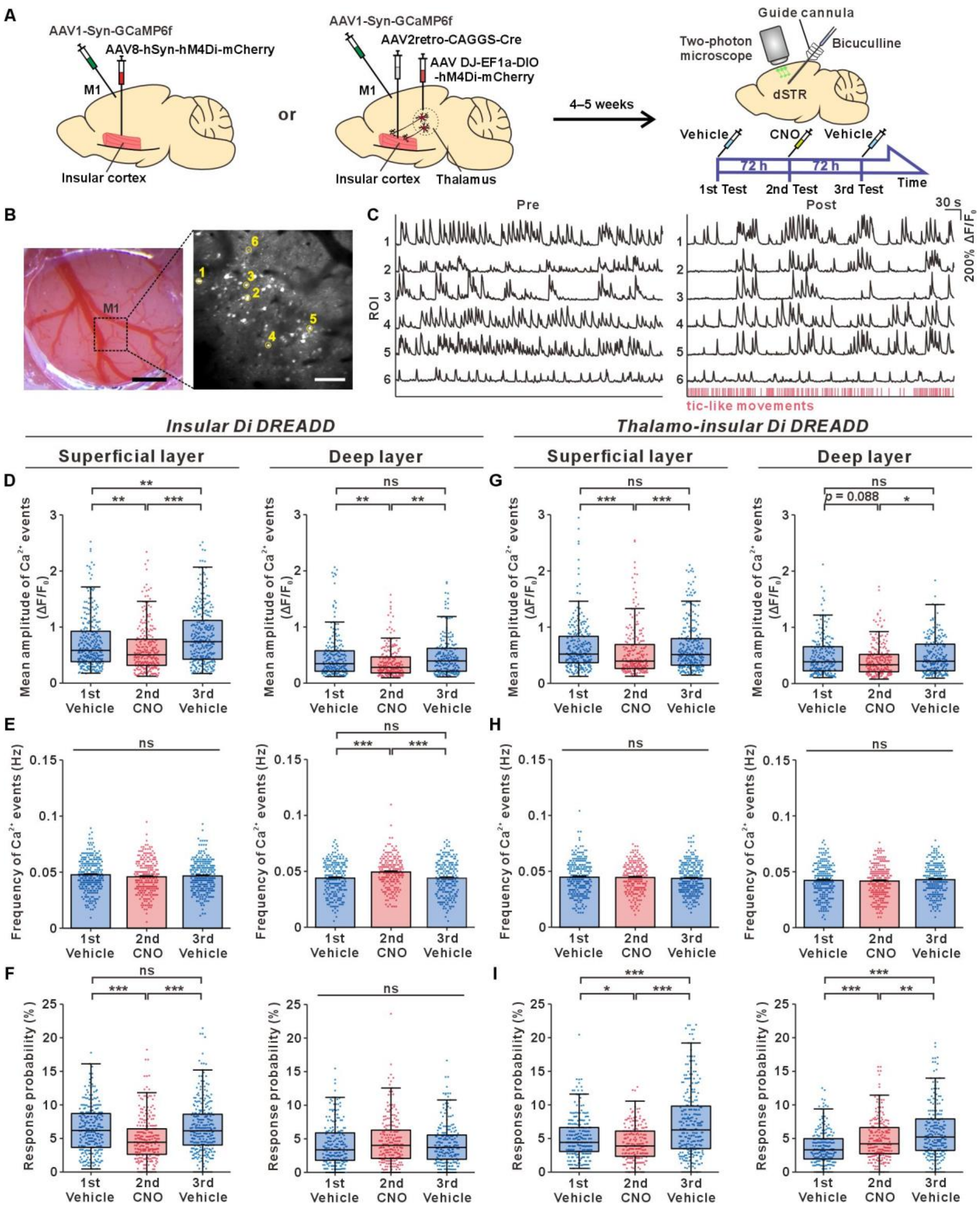


**Figure S4. Chemogenetic inhibition of IC or the dmTh-IC pathway alters M1 activity in a layer-specific manner. Related to Figure 6.**

(A) Schematic of the experimental setup for two-photon $Ca^{2+}$ imaging during chemogenetic inhibition of IC or of dmTh neurons projecting to IC. To inhibit IC, AAV8-hSyn-hM4Di-mCherry was bilaterally injected into three rostrocaudally distinct IC sites. To specifically inhibit the dmTh-IC pathway, AAV2retro-CAGGS-Cre was bilaterally injected into IC, and AAV-DJ-EF1α-DIO-hM4Di-mCherry into bilateral dmTh. AAV1-Syn-GCaMP6f was injected into the ipsilateral M1. Tic-like movements were induced by unilateral bicuculline injections into the dorsal striatum (dSTR) under vehicle (1st test), CNO (10 mg/kg; 2nd test), and vehicle (3rd test) conditions.

(B) Left, cranial window over M1. Scale bar, 500 μm. Right, representative two-photon image of GCaMP6f-expressing neurons in the superficial layer of M1; yellow circles indicate detected neurons. Scale bar, 100 μm.

(C) Representative $Ca^{2+}$ signals from six detected neurons in (B) during the pre- (left) and post- (right) injection periods. Tic events are indicated by red bars. Synchronized tic-associated M1 activity was observed.

(D-F) Effect of IC inhibition on $Ca^{2+}$ activity in the superficial (layer 2/3; n = 286, 250, and 298 neurons for the 1st, 2nd, and 3rd tests, respectively; n = 5 mice) and deep (layer 5/6; n = 216, 215, and 212 neurons for the three sessions, respectively; n = 5 mice) layers of the ipsilateral M1. IC inhibition reduced $Ca^{2+}$ event amplitudes in both layers (D). In the superficial layer, event frequency was unaffected, while in the deep layer it was significantly increased (E). Response probability was significantly reduced in the superficial layer, but not in the deep layer (F).

(G-I) Effect of the dmTh-IC pathway inhibition on $Ca^{2+}$ activity in the superficial (n = 260, 223, and 275 neurons for the three sessions, respectively; n = 5 mice) and deep (n = 214, 230, and 222 neurons for the three sessions, respectively; n = 5 mice) layers of the ipsilateral M1. Pathway-specific inhibition reduced event amplitudes in the superficial layer, but not in the deep layer (G). The same manipulation did not affect event frequency (H), but modulated response probability in opposite directions in the two layers (I).

(D-I) Bar graphs show mean ± SEM. Box plots indicate the median, 25th, and 75th percentiles, with whiskers representing data within a 1.5 × interquartile range. Each dot represents an individual neuron. Statistical analyses were performed using the Kruskal-Wallis test followed by Dunn's multiple comparison test with Holm correction (D, F, G, and I) or one-way ANOVA followed by pairwise *t*-tests with Holm correction (E and H). ns, not significant; *p < 0.05; **p < 0.01; ***p < 0.001.

See also Video S3.

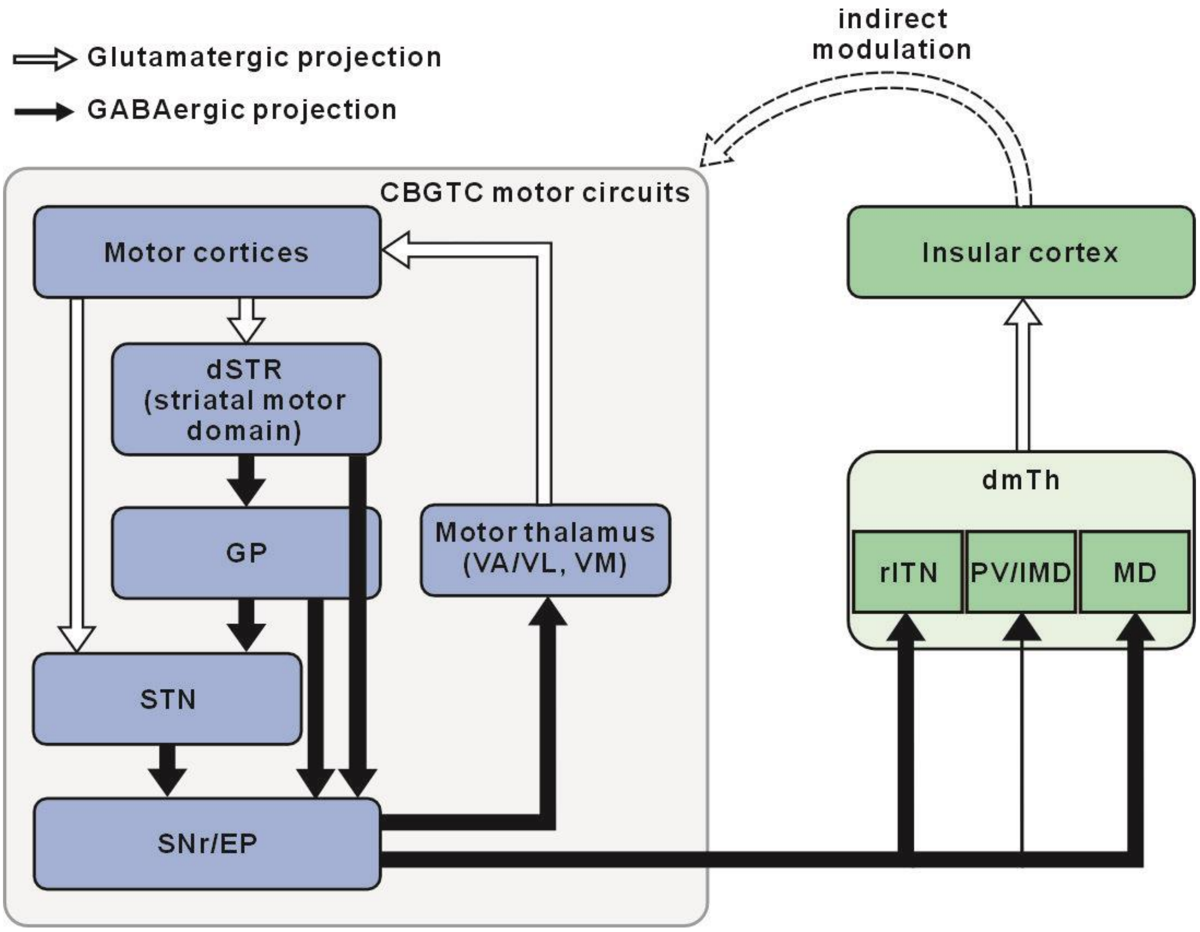


**Figure S5. Motor-limbic crosstalk underlying tic generation. Related to Figure 6.**

Schematic of a “motor-to-limbic” open loop within the CBGTC circuits. This pathway illustrates how abnormal motor signals can propagate into limbic structures such as the insular cortex, thereby contributing to tic generation. STN, subthalamic nucleus.

**Video S1. Tic-like movements and tic-related EEG spikes induced by striatal bicuculline injection. Related to Figure 1.**

Representative video showing tic-like movements and EEG recordings from M1. Bicuculline was unilaterally injected into the left dorsal striatum, inducing tic-like movements primarily in the contralateral forelimb and orofacial (mouth corner) regions. Colored dots indicate the positions of user-defined body parts tracked using DeepLabCut. In the EEG trace (bottom), the red vertical line marks the time point corresponding to the behavioral video.

**Video S2. Suppression of motor tics by chemogenetic inhibition of IC or the dmTh-IC pathway. Related to Figure 5.**

Representative video showing the suppression of motor tics by chemogenetic inhibition of IC or the dmTh-IC pathway. The video consists of four clips: the first two show behavior without and with CNO administration during IC inhibition, and the latter two show behavior without and with CNO administration during the dmTh-IC pathway inhibition. All clips were recorded ~15 min after bicuculline injection, when tic-like movements were most prominent.

**Video S3. Synchronized neuronal activity in M1 following striatal bicuculline injection. Related to Figure 6.**

Representative video showing $Ca^{2+}$ signals of multiple neurons in the ipsilateral M1 before and after striatal bicuculline injection, recorded using two-photon microscopy. Left: pre-injection; right: post-injection. Playback speed, 7x.
